# Leaf ionomic stability shapes drought adaptation in pearl millet

**DOI:** 10.64898/2026.11.12.766145

**Authors:** Princia Nakombo-Gbassault, Sebastian Arenas, Pablo Affortit, Awa Faye, Paulina Flis, Bassirou Sine, Daniel Moukouanga, Pascal Gantet, Ephrem Kosh-Komba, Ndjido Kane, Malcolm Bennett, Darren Wells, Philippe Cubry, Elizabeth Bailey, Vincent Segura, Yves Vigouroux, Laurent Laplaze, Alexandre Grondin

## Abstract

Pearl millet is a staple cereal of semi-arid West Africa, where early-season water deficit constrains vegetative establishment and yield. While the genetic architecture of the pearl millet foliar ionome has been characterised under well-watered conditions, how this ionome reorganises under drought, and whether this reorganisation has a distinct genetic basis, remains unknown. We profiled the drought-response of 17 foliar elements in 175 diverse pearl millet inbred lines over two seasons in Senegal. The foliar concentration of two co-regulated groups of elements changed reproducibly under early water stress across both years. Leaf ion stability, rather than plasticity, was associated with the maintenance of flowering time and plant biomass, suggesting that drought adaptation in pearl millet relies more on ionomic stability than on adaptive shifts in elemental composition. Among the 17 elements, only potassium showed a significant genotype-by-treatment interaction suggesting that genetic variation contributes to differences in potassium response to drought. Genome-wide association mapping on potassium plasticity in response to drought identified a locus, designated qKP-6, that limits potassium depletion under stress without penalising constitutive potassium levels. This locus represents a candidate region for the genetic control of potassium plasticity under drought. This work provides new insights into the genetic control of mineral nutrition under water deficit in pearl millet, and highlights leaf potassium concentration as a genetically controlled component of drought-responsive ionomic variation with potential implications for drought tolerance.

## Introduction

Water scarcity, exacerbated by climate change, poses a major threat to the sustainability of agricultural production in arid and semi-arid regions where rainfall is highly variable and drought events are projected to increase in both frequency and severity (Intergovernmental Panel On Climate Change (IPCC), 2023; Rezaei *et al*., 2023). In these predominantly rainfed systems, food security critically depends on drought-tolerant crops capable of sustaining at least minimal but reliable yields under water-limited conditions (Satyavathi *et al*., 2021).

Among drought-tolerant crops, pearl millet (*Pennisetum glaucum (L.) R. Br.*), a C4 cereal domesticated in the Sahel, stands out for its exceptional resilience to both water and heat deficits (Debieu *et al*., 2017). It is widely cultivated in regions receiving as little as 300 to 500 mm of annual rainfall, where only few other cereal species can survive (Bidinger *et al*., 1987; Pucher *et al*., 2015). This resilience relies on a diversity of adaptive strategies, including tolerance, avoidance and escape mechanisms, operating at different developmental stages from vegetative growth to flowering and grain filling (Shrestha *et al*., 2023). Recent crop-modelling evidence indicates that early drought stress, occurring before or during panicle initiation, is the most damaging water-stress pattern for pearl millet in the Sahel, causing grain yield losses of up to 44% and biomass reductions of approximately 40% (De la Fuente *et al*., 2024). Climate-based simulations further show that early drought stress occurs more frequently than terminal stress for pearl millet in this region, a trend expected to be reinforced in the context of changing climate, underscoring its critical impact on crop establishment in rainfed systems (De la Fuente *et al*., 2024). Because this stress coincides with vegetative development, a period of rapid mineral uptake and partitioning (Marschner and Marschner, 2012), it may have particularly pronounced effects on the leaf ionome, defined as the composition of plant tissue in mineral nutrients and trace elements (Salt *et al*., 2008; Srivastava *et al*., 2022), with potential implications for photosynthesis, osmotic adjustment, and stress signalling (Farooq *et al*., 2009).

Water availability is a major determinant of plant growth and mineral nutrition. Beyond its direct effects on cell expansion and photosynthesis, reduced water availability profoundly reshapes plant mineral composition and regulation (D’Oria *et al*., 2022). Nutrient responses to drought are no longer viewed as a passive consequence of reduced transpiration or growth alone, but as actively regulated processes involving specific membrane transporters, signalling pathways, and metabolic adjustments (Bisht *et al*., 2026). Among essential macronutrients, potassium (K), calcium (Ca), and magnesium (Mg) play central but contrasting roles under water limitation: K contributes to osmotic adjustment and stomatal regulation, Ca functions as a key secondary messenger in stress signalling, and Mg deficiency reduces whole-plant water-use efficiency (Tränkner *et al*., 2016; Johnson *et al*., 2022; Liu *et al*., 2022). Under water deficit, studies have revealed reproducible shifts in elemental profiles, including changes in the concentrations of Na, K, Mg, Mn or Cu, whose direction and magnitude depend on stress intensity, duration, and species (D’Oria *et al*., 2022; Quagliata *et al*., 2025). Importantly, these changes can occur prior to the appearance of visible morphological symptoms, indicating that the ionome is an early and sensitive indicator of drought responses (Danku *et al*., 2013; D’Oria *et al*., 2022). Thus, drought-induced changes in elemental composition can offer an integrative view of physiological adjustment to water limitation, while genotypic variation in the magnitude of these changes may reflect ionomic plasticity. The differential regulation of these elements under drought therefore reflects an integrated reprogramming of the plant ionome, which may influence stress tolerance and ultimately yield.

Ionomics, the high-throughput profiling of elemental composition across genotypes and environments, provides an integrative framework to investigate the genetic determinants of plant responses to environmental constraints (Salt *et al*., 2008; White and Broadley, 2009; Bouain *et al*., 2019). Yet, the genetic bases of ionomic variation and plasticity under water-limited conditions remain incompletely characterized, particularly in crops adapted to marginal environments. Genome-wide association studies of the ionome have been conducted primarily under near-optimal conditions, limiting our understanding of ionomic plasticity under drought and the detection of genotype × environment interactions in response to this stress. In pearl millet specifically, the genetic architecture of foliar mineral variation and responsiveness under water deficit remains unexplored, leaving unresolved to what extent water stress-induced plasticity is under genetic control and how it contributes to plant performance under water limitation.

Here, we combined field-based ionomic phenotyping with genome-wide association analyses in a diverse pearl millet panel evaluated under two contrasting water regimes over two years to identify the genomic regions associated with constitutive foliar elemental variation and its plastic response to water deficit. We hypothesised that 1) the leaf ionomic profiles under drought would exhibit substantial genotypic variation in pearl millet and be associated with plant performance, and 2) ion content are under genetic control, with loci underlying constitutive ionomic content being largely distinct from those controlling ionomic plasticity under drought.

## Materials and Methods

### Plant material

A total of 175 pearl millet (*Pennisetum glaucum* (L.) R. Br.) inbred lines were selected from the Pearl Millet Genetic Association Panel (PMiGAP; Sehgal *et al*., 2015; Varshney *et al*., 2017), which captures the genetic diversity of the species from Africa and India. Detailed composition of the lines used here is provided in Nakombo-Gbassault *et al*. (2025).

### Field trials and experimental design

Field trials were conducted at the National Center for Agronomic Research (CNRA, ISRA, Bambey, Senegal; 14°42’N, 16°28’W) during the hot dry off-seasons of 2021 and 2022 on a loamy sand soil (84.8% sand, 7.9% silt, 7.2% clay; Diongue *et al*., 2022). An alpha-lattice design was used with two water regimes: well-watered (WW) and vegetative-stage water stress (WS), with four replicates per treatment. In both years, prior to planting, soil samples were collected from thirteen locations distributed across the four replicate blocks, at four depth intervals (0-20, 20-60, 60-100, and 100-140 cm).

Since no rainfall occurs during this period of the year, water stress was imposed simply by withholding irrigation from 21 to 49 days after sowing (DAS) in 2021 and from 21 to 42 DAS in 2022, corresponding to stress durations of 28 and 21 days respectively. These conditions defined four environments used in subsequent analyses: WW21 (well-watered) and WS21 (water-stress) for the 2021 trial, and WW22 and WS22 for the 2022 trial. Relative soil water content was monitored from 0 to 160 cm using DIVINER probes spread across both treatments.

At the end of the stress period, three plants per plot were sampled for leaf ionomic profiling, root anatomical phenotyping, and aboveground biomass measurement, as described in Nakombo-Gbassault *et al*. (2025). For ionomic analysis, the last fully ligulated leaf from the main tiller of each harvested plant was collected, washed, dried, and pooled per plot before digestion and quantification by inductively coupled plasma mass spectrometry (ICP-MS) as described (Nakombo-Gbassault *et al*., 2025). Foliar concentrations of 17 elements were quantified, including macro- and micronutrients: potassium (K), sodium (Na), magnesium (Mg), calcium (Ca), phosphorus (P), sulfur (S), iron (Fe), zinc (Zn), copper (Cu), cadmium (Cd), nickel (Ni), manganese (Mn), molybdenum (Mo), cobalt (Co), lithium (Li), rubidium (Rb), and strontium (Sr). Plant available soil elements were also quantified using 1M NH_4C_O_3-_DTPA extraction. For root anatomy, crown root segments were sampled and analysed by laser ablation tomography to quantify stele area, total metaxylem vessel area and number, sclerenchyma proportion, and overall root cross-sectional area, as described in Affortit *et al*. (2024).

After the stress period, plants were re-irrigated until maturity, and phenology as well as yield and yield-component traits were recorded, including plant height, shoot biomass at maturity, spike number, grain weight per plant and harvest index, following procedures described in Affortit *et al*., (2024). Trait list and acronyms are described in Supplementary Table S1.

### Stress response analyses

For each phenotypic trait including element concentrations, root anatomical and agromorphological traits, best linear unbiased estimates (BLUEs) and broad-sense heritability were estimated for each treatment-year combination using the SpATS spatial model (Rodríguez-Álvarez *et al*., 2018), implemented in the statgenSTA R package (Van Rossum, 2020). The model was fitted to data from a resolvable incomplete block design. Genotype was fitted as a fixed effect to obtain BLUEs and as a random effect to estimate broad-sense heritability using the generalised heritability approach described by Oakey *et al*., (2006).

Two complementary stress indices were computed per trait and genotype, restricted to genotypes present in both water regimes within the same year:

i. the plasticity index, expressed as a log2 ratio (LR):

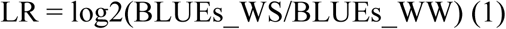

quantifies relative plasticity, with LR = 0 indicating no change, negative values indicating stress-induced depletion, and positive values indicating accumulation under stress;

(ii) the Stress Tolerance Index (STI; Fernandez, 1992):

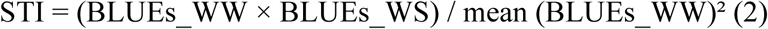

captures absolute performance across both water regimes. Both indices were computed for ionomic and agromorphological traits.

To identify elements whose response to early water deficit is repeatable across seasons, the direction and magnitude of each element’s response were characterised per year by the percentage change in mean concentration and the mean log2 ratio (LR) across genotypes. WW and WS BLUEs of the same genotypes were compared with a paired t-test when residuals were normally distributed (Shapiro-Wilk *p* > 0.05), and with a Wilcoxon signed-rank test otherwise. An element response was considered robust when significant in both years with the same direction of change.

Because soil sampling preceded treatment imposition, it reflects baseline spatial heterogeneity rather than treatment-induced variation in soil ion availability. Multivariate correspondence between soil (0-20 cm layer, ICP-MS values) and foliar ion concentrations was tested separately for each of the four environments using a Mantel test on the 15 elements common to both datasets (standardised Euclidean distances, 999 permutations; Mantel, 1967).

### Multivariate analysis of the foliar ionome

Principal component analysis (PCA) was performed on foliar ion concentrations (BLUEs) to characterise the variation of the 17 elements. A global PCA was first conducted using data from all four environments (WW21, WS21, WW22, and WS22) to characterise the overall structure of ionomic variation. To isolate the treatment-driven component of variation from inter-annual differences, separate PCAs were then performed for each year using the corresponding WW and WS samples. For these year-specific analyses only, the sign of the principal components was reversed when necessary, so that water-stressed genotypes consistently projected towards negative scores on the primary stress-response axis, facilitating direct comparison of this axis between years. For each configuration, the contribution of each element to each component (percentage of the component’s total inertia) and the element loading (coordinate on the component axis) were reported to identify the principal drivers of the stress axis. For the stress-response axis, scores under each treatment (PC_WW, PC_WS) and their difference (ΔPC = PC_WS - PC_WW) were computed per genotype and year.

### Correlation analyses

Pairwise Pearson correlations among the 17 foliar elements were computed within each environment on BLUEs and their significance assessed with a 5% false discovery rate (FDR) correction (Benjamini and Hochberg, 1995). Pairs were classified as constitutive (significant with consistent sign in all four environments), never significant, or context dependent. Conservation of the correlation structure between water regimes was tested per year using Mantel tests (Mantel, 1967) and its significance tested using 999 permutations. Hierarchical clustering (average linkage) was applied to the mean correlation matrix across the four environments to identify co-regulation modules of ions. Ions within each module were then combined into a composite score, which was used in subsequent analyses in place of the individual ions.

Correlations between the ionome and agromorphological traits were assessed at three levels: individual elements (mono-ion), modules, and the combined-PCA stress-response axis (multi-ion). Two complementary approaches were applied at each level: (i) within-environment Pearson correlations, retaining associations significant in both years with a consistent sign; and (ii) linear regression pooling both years, with year as a fixed covariate and cluster-robust standard errors at the genotype level (vcovCL, Zeileis *et al*., 2020). FDR correction at 5% was applied within each trait category, with STI and LR indices corrected independently. For all pooled models, a predictor x Year interaction term additionally tested whether association strength differed between years.

### Predicting genotype-level drought tolerance from the constitutive ionome

To predict genotype-level drought tolerance, the STI of shoot biomass and of grain weight were used. Three predictors were compared. The first approach used a simple linear regression to assess the well-watered score of the PC2 axis as a predictor of the STI. The second approach used a linear regression to identify the ion most strongly correlated with the STI as a predictor. The third approach used a penalized regression (LASSO; Tibshirani, 1996; glmnet package, Friedman et al., 2010) on all measured elements to identify a multi-ion signature predictor, with λ chosen at the one-standard-error rule and with an ion selection frequency over 20 refits. In the three approaches, the predictive accuracy, namely the correlation between observed and out-of-fold predicted values, was estimated by repeated genotype-grouped 5-fold cross-validation (20 repetitions) with identical partitions for all predictors, and predictors were compared using bootstrap intervals of the differences (10,000 resamples of genotypes). All predictors were evaluated on the same held-out genotypes, so that accuracy reflects prediction of new genotypes rather than model fit.

The predicted score (centred on the year mean) for each STI, calculated as the mean out-of-fold for the best predictor (highest mean predictive accuracy) of the three approaches, was used to classify genotypes as tolerant, intermediate or sensitive using tercile thresholds, as was the observed STI. To verify that predicted scores separated genotypes that were tolerant from those that were sensitive for the observed STI, the scores of these two groups were compared with a Wilcoxon test. Fisher’s exact and Spearman tests were used to test whether genotypic classification between STI for shoot biomass and STI for grain weight was conserved, on predicted and observed values. To test if population structure was a potential confounder, a PCA using the genotypic data from the panel was performed and the first two components were selected. The first two components were correlated with the predicted scores and included as covariates in the regression of the observed STI on the predicted score.

### Estimation of phenotypic variance components

To partition the variation of each principal component and each element into constitutive and condition-specific genetic components, a linear mixed model was fitted to genotype BLUEs by restricted maximum likelihood (REML, lme4; Bates *et al*., 2015):

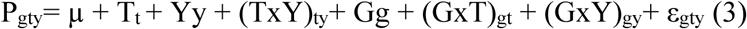

where T and Y are fixed effects of treatment and year, (T×Y) their fixed interaction, G the random effect of genotype, and G×T and G×Y the random genotype-by-treatment and genotype-by-year interactions. Year was treated as fixed because it has only two levels. Random effects were tested by likelihood-ratio tests comparing the full model to the model without the effect (with statistical threshold *p* < 0.05) and, for principal components, fixed effects by Type II F-tests (Satterthwaite approximation, lmerTest, Kuznetsova *et al*., 2017). For elements, each random component is expressed as a percentage of the total random variance. For principal components, the variance of each fixed term (sum-to-zero coding) was added to the random components so that the treatment effect could be compared with the genetic components (Nakagawa and Schielzeth, 2013). The principal component with the highest treatment percentage was retained as the stress-response axis. In the year-specific PCAs, the treatment share was estimated as η² = SS_T/SS_total from the linear model:

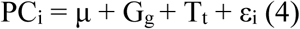

with Genotype (G) and Treatment (T) as fixed effect.

### Genome-wide association analysis (GWAS)

A total of 269,848 SNPs were obtained from whole-genome sequencing of 147 inbred lines from the PMiGAP panel, after filtering for minor allele frequency (MAF > 0.05) and missing data rate (< 20% per SNP and per genotype). Procedures for filtering, imputation, and population structure inference are described in Nakombo-Gbassault *et al*., (2025).

GWAS was performed on ion BLUEs for each of the four environments separately (WW21, WW22, WS21, WS22), using four complementary models correcting for population structure and/or genetic relatedness through different approaches: LFMM2 (Frichot *et al*., 2014), BLINK (Huang *et al*., 2019), EMMA (Kang *et al*., 2008) and CMLM (Zhang *et al*., 2010). For ions showing a significant genotype × treatment interaction, GWAS was performed on the corresponding plasticity index (LR) to identify loci associated with variation in the response to water deficit.

### Multi-environment meta-analysis of GWAS

For each of the four GWAS models, results across environments were integrated independently using the fixed-effect (*Fe*) procedure of the metaGE framework in R (De Walsche *et al*., 2025). Because the same genotypes were evaluated across environments, the correlation between environment-specific GWAS was first estimated to account for this non-independence in the model.

In metaGE, the estimated effect (β) and *p*-value of each marker from each environment-specific GWAS is converted into a signed Z-score:

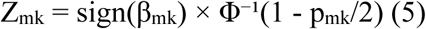

where Z_mk i_s the Z-score of marker m in environment k, β_mk a_nd p_mk i_ts estimated effect and *p*-value, and Φ⁻¹ the quantile function of the standard normal distribution.

Two complementary tests were applied: (i) a model without treatment contrast, identifying markers with effects consistent across all environments, and (ii) a model with an explicit contrast opposing irrigation and water deficit, testing for marker effects within each water regime and differences between regimes. The no-contrast model was applied (i) across all four environments, (ii) within each water regime separately (WW21 and WW22 combined or WS21 and WS22 combined), and (iii) on the plasticity index (LR, both years combined). The contrast model was applied across all four environments. Significance was set at *p* < 10^⁻5^. To delimit a region of interest around each significant SNP, pairwise linkage disequilibrium (r²) was calculated using PLINK v1.9 (Chang *et al*., 2015) within a ± 50 kb window centred on the significant marker. SNPs with r² > 0.5 were considered genetically linked and grouped with the significant SNP to define the boundaries of the associated region. Candidate genes within this region were identified using the functional annotation of the pearl millet reference genome (ASM217483v2; (Varshney *et al*., 2017).

### Allelic class effect analysis

For each significant marker, trait associations were tested across the three allelic classes (homozygous reference, heterozygous, homozygous alternative) determined from VCF genotype calls, using linear mixed models fitted by maximum likelihood (REML = FALSE), which allows likelihood ratio tests between models differing in fixed effects:

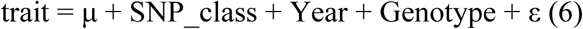

where SNP_class and Year are fixed effects, Genotype is a random effect of line identity accounting for repeated measurements across years, and ε is the residual. Significance was assessed by likelihood ratio test against a null model without the SNP_class term, and effect size was estimated by the increase in marginal R² relative to this null model (ΔR²m; Nakagawa and Schielzeth, 2013). Estimated marginal means of allelic classes were compared using Tukey-adjusted pairwise contrasts. Dependence of the allelic effect on water regime was tested using a SNP class × treatment interaction:

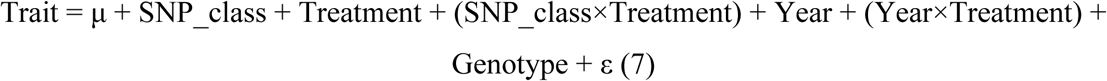

To assess whether the SNP effect was independent of the phenological variation, the association between each trait and flowering time under well-watered conditions (DTF_WW) was first tested by replacing SNP_class with DTF_WW in model (6); DTF_WW was then included as a covariate in both the full and null models:

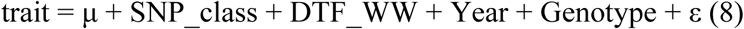

and the change in the difference between homozygous classes was expressed as a percentage reduction.

## Result

### Genotypic and interannual variation in the foliar ionomic response to early water deficit

To characterise the foliar ionomic response to early water stress, we quantified 17 mineral elements in leaves of 175 pearl millet inbred lines evaluated under well-watered (WW) and water-stressed (WS) conditions over two seasons (2021 and 2022) at Bambey (Senegal). Under water stress, the 17 measured elements varied widely among genotypes, with coefficients of variation ranging from 12.4% (S) to 44.8% (Na) in 2021 and from 12.4% (S) to 51.9% (Li) in 2022 (Table 1). Broad-sense heritability also showed substantial variation, from low for lithium (H² = 0.01 in 2022) and nickel (H² = 0.04 in 2022) to high for cadmium (H² = 0.77-0.84) and cobalt (H² = 0.80-0.82) indicating strong differences in the genetic control of ion accumulation in pearl millet (Table 1). Soil ion concentrations were sampled from thirteen locations spread across the four replicate blocks, at four depths per location. These soil values did not match mean foliar ion concentrations in any of the four environments (Mantel tests, *r* = −0.37 to 0.09, all *p* > 0.45), suggesting that differences between genotypes were independent of differences in soil ion contents (Supplementary Table S2).

**Table 1:**
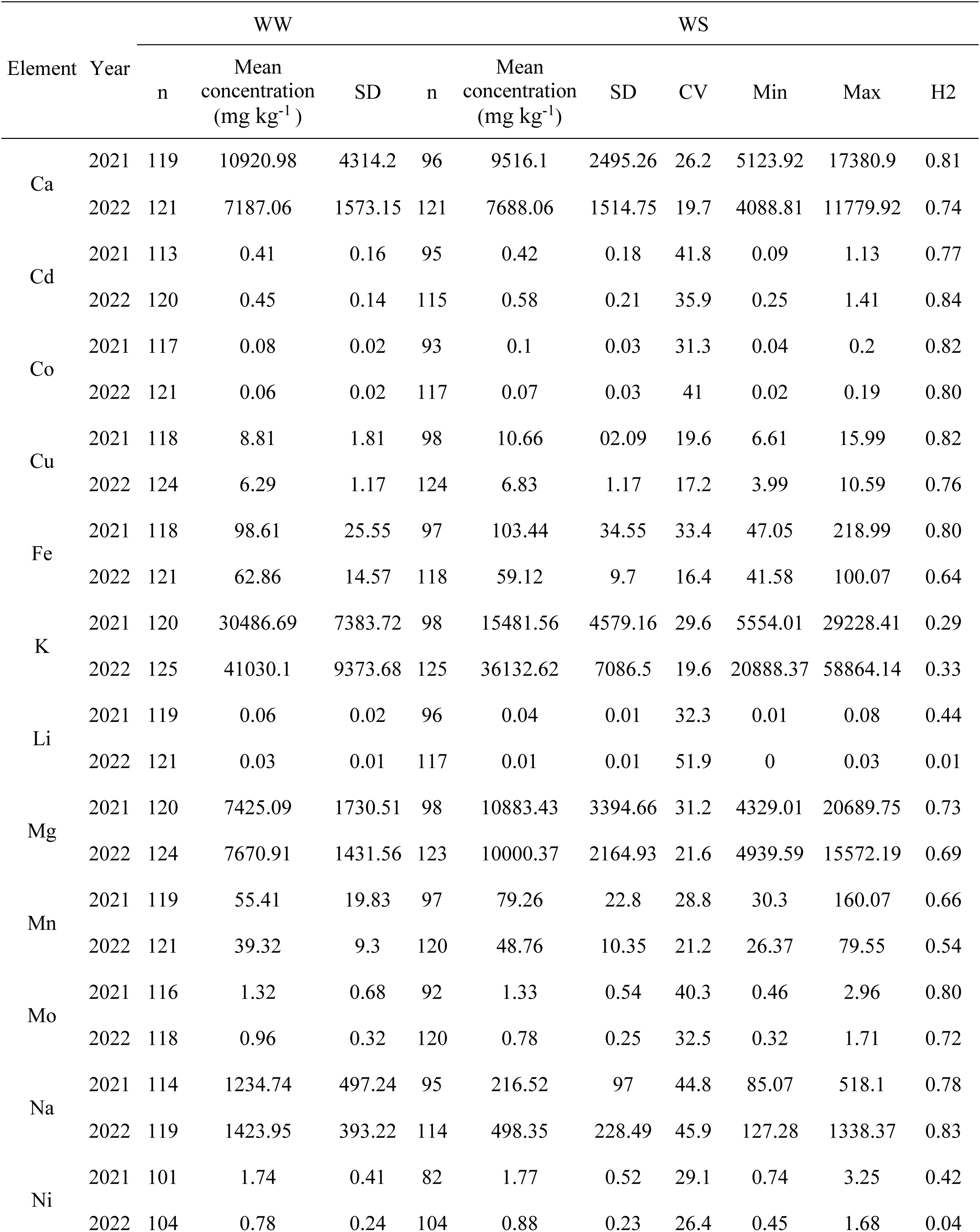
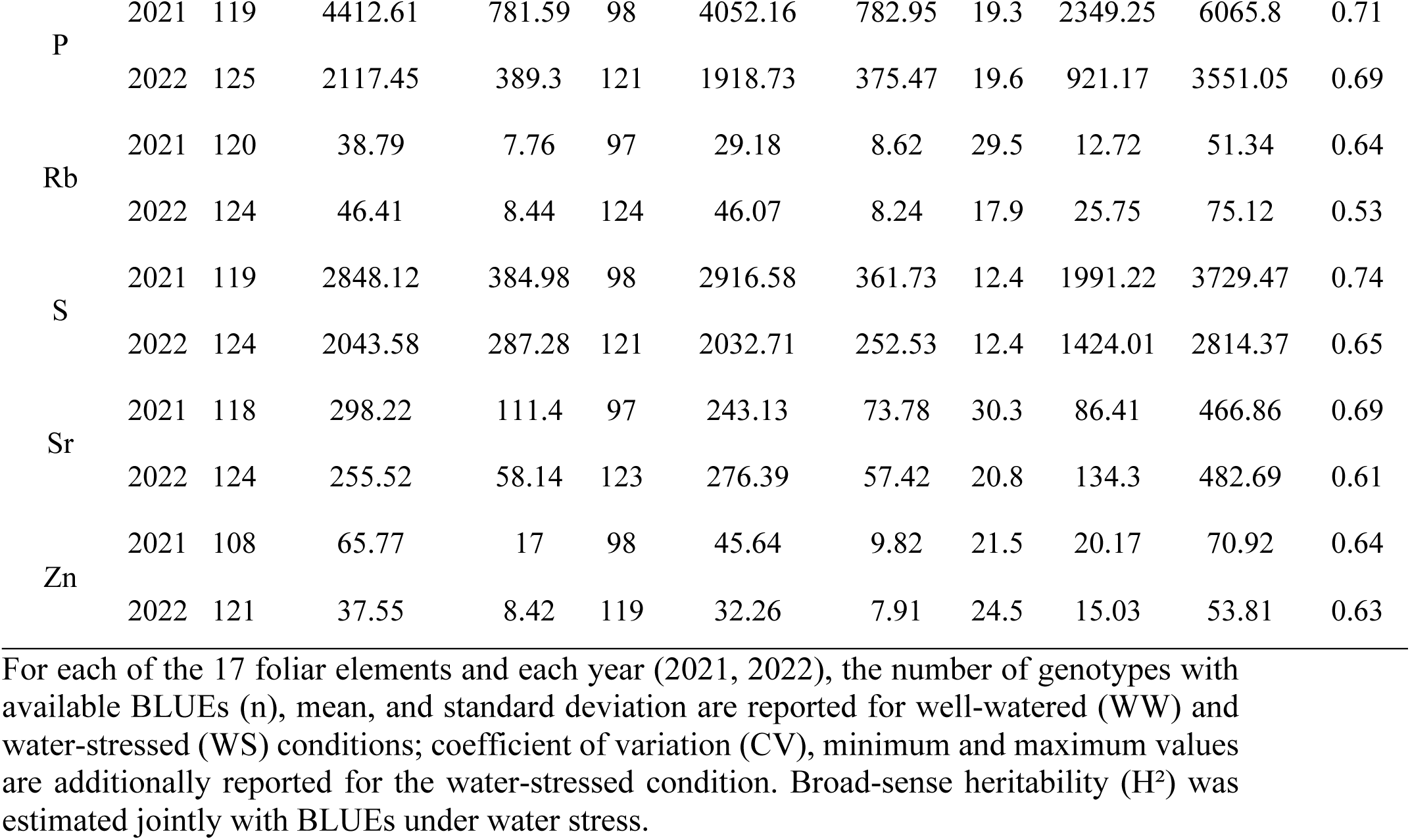
Descriptive statistics and broad-sense heritability of foliar ion concentrations under well-watered and water-stressed conditions.

| Element | Year | WW |  |  | WS |  |  |  |  |  |
|---|---|---|---|---|---|---|---|---|---|---|
|  |  | n | Mean concentration (mg kg<sup>-1</sup>) | SD | n | Mean concentration (mg kg<sup>-1</sup>) | SD | CV | Min | Max | H2 |
| Ca | 2021 | 119 | 10920.98 | 4314.2 | 96 | 9516.1 | 2495.26 | 26.2 | 5123.92 | 17380.9 | 0.81 |
|  | 2022 | 121 | 7187.06 | 1573.15 | 121 | 7688.06 | 1514.75 | 19.7 | 4088.81 | 11779.92 | 0.74 |
| Cd | 2021 | 113 | 0.41 | 0.16 | 95 | 0.42 | 0.18 | 41.8 | 0.09 | 1.13 | 0.77 |
|  | 2022 | 120 | 0.45 | 0.14 | 115 | 0.58 | 0.21 | 35.9 | 0.25 | 1.41 | 0.84 |
| Co | 2021 | 117 | 0.08 | 0.02 | 93 | 0.1 | 0.03 | 31.3 | 0.04 | 0.2 | 0.82 |
|  | 2022 | 121 | 0.06 | 0.02 | 117 | 0.07 | 0.03 | 41 | 0.02 | 0.19 | 0.80 |
| Cu | 2021 | 118 | 8.81 | 1.81 | 98 | 10.66 | 02.09 | 19.6 | 6.61 | 15.99 | 0.82 |
|  | 2022 | 124 | 6.29 | 1.17 | 124 | 6.83 | 1.17 | 17.2 | 3.99 | 10.59 | 0.76 |
| Fe | 2021 | 118 | 98.61 | 25.55 | 97 | 103.44 | 34.55 | 33.4 | 47.05 | 218.99 | 0.80 |
|  | 2022 | 121 | 62.86 | 14.57 | 118 | 59.12 | 9.7 | 16.4 | 41.58 | 100.07 | 0.64 |
| K | 2021 | 120 | 30486.69 | 7383.72 | 98 | 15481.56 | 4579.16 | 29.6 | 5554.01 | 29228.41 | 0.29 |
|  | 2022 | 125 | 41030.1 | 9373.68 | 125 | 36132.62 | 7086.5 | 19.6 | 20888.37 | 58864.14 | 0.33 |
| Li | 2021 | 119 | 0.06 | 0.02 | 96 | 0.04 | 0.01 | 32.3 | 0.01 | 0.08 | 0.44 |
|  | 2022 | 121 | 0.03 | 0.01 | 117 | 0.01 | 0.01 | 51.9 | 0 | 0.03 | 0.01 |
| Mg | 2021 | 120 | 7425.09 | 1730.51 | 98 | 10883.43 | 3394.66 | 31.2 | 4329.01 | 20689.75 | 0.73 |
|  | 2022 | 124 | 7670.91 | 1431.56 | 123 | 10000.37 | 2164.93 | 21.6 | 4939.59 | 15572.19 | 0.69 |
| Mn | 2021 | 119 | 55.41 | 19.83 | 97 | 79.26 | 22.8 | 28.8 | 30.3 | 160.07 | 0.66 |
|  | 2022 | 121 | 39.32 | 9.3 | 120 | 48.76 | 10.35 | 21.2 | 26.37 | 79.55 | 0.54 |
| Mo | 2021 | 116 | 1.32 | 0.68 | 92 | 1.33 | 0.54 | 40.3 | 0.46 | 2.96 | 0.80 |
|  | 2022 | 118 | 0.96 | 0.32 | 120 | 0.78 | 0.25 | 32.5 | 0.32 | 1.71 | 0.72 |
| Na | 2021 | 114 | 1234.74 | 497.24 | 95 | 216.52 | 97 | 44.8 | 85.07 | 518.1 | 0.78 |
|  | 2022 | 119 | 1423.95 | 393.22 | 114 | 498.35 | 228.49 | 45.9 | 127.28 | 1338.37 | 0.83 |
| Ni | 2021 | 101 | 1.74 | 0.41 | 82 | 1.77 | 0.52 | 29.1 | 0.74 | 3.25 | 0.42 |
|  | 2022 | 104 | 0.78 | 0.24 | 104 | 0.88 | 0.23 | 26.4 | 0.45 | 1.68 | 0.04 |
| P | 2021 | 119 | 4412.61 | 781.59 | 98 | 4052.16 | 782.95 | 19.3 | 2349.25 | 6065.8 | 0.71 |
|  | 2022 | 125 | 2117.45 | 389.3 | 121 | 1918.73 | 375.47 | 19.6 | 921.17 | 3551.05 | 0.69 |
| Rb | 2021 | 120 | 38.79 | 7.76 | 97 | 29.18 | 8.62 | 29.5 | 12.72 | 51.34 | 0.64 |
|  | 2022 | 124 | 46.41 | 8.44 | 124 | 46.07 | 8.24 | 17.9 | 25.75 | 75.12 | 0.53 |
| S | 2021 | 119 | 2848.12 | 384.98 | 98 | 2916.58 | 361.73 | 12.4 | 1991.22 | 3729.47 | 0.74 |
|  | 2022 | 124 | 2043.58 | 287.28 | 121 | 2032.71 | 252.53 | 12.4 | 1424.01 | 2814.37 | 0.65 |
| Sr | 2021 | 118 | 298.22 | 111.4 | 97 | 243.13 | 73.78 | 30.3 | 86.41 | 466.86 | 0.69 |
|  | 2022 | 124 | 255.52 | 58.14 | 123 | 276.39 | 57.42 | 20.8 | 134.3 | 482.69 | 0.61 |
| Zn | 2021 | 108 | 65.77 | 17 | 98 | 45.64 | 9.82 | 21.5 | 20.17 | 70.92 | 0.64 |
|  | 2022 | 121 | 37.55 | 8.42 | 119 | 32.26 | 7.91 | 24.5 | 15.03 | 53.81 | 0.63 |

For each of the 17 foliar elements and each year (2021, 2022), the number of genotypes with available BLUEs (n), mean, and standard deviation are reported for well-watered (WW) and water-stressed (WS) conditions; coefficient of variation (CV), minimum and maximum values are additionally reported for the water-stressed condition. Broad-sense heritability (H²) was estimated jointly with BLUEs under water stress.

The ionomic profile was more strongly disturbed in response to vegetative drought stress in 2021 than in 2022, with a mean plasticity index (LR) across the 17 elements of −0.23 in 2021 versus −0.12 in 2022. Comparing the two water regimes revealed three clear and repeatable patterns of ionomic response to vegetative drought (Fig. 1). A first group of four elements increased significantly under water deficit in both years. This group included magnesium (Mg), cobalt (Co), manganese (Mn), and copper (Cu). A second group of five elements decreased significantly under water deficit in both years. This group comprised sodium (Na), potassium (K), zinc (Zn), lithium (Li), and phosphorus (P). A third group, made up of calcium and strontium, reversed direction between years: both elements decreased under water deficit in 2021 but increased in 2022. The treatment-by-year interaction was significant for 14 of the 17 elements (Supplementary Table S3). Lithium, sodium and nickel were the only elements with no significant interaction. Together with the moderate-to-high heritability of several elements, these results indicate that the early ionomic response to water deficit in pearl millet is reproducible and structured.

**Fig. 1.**
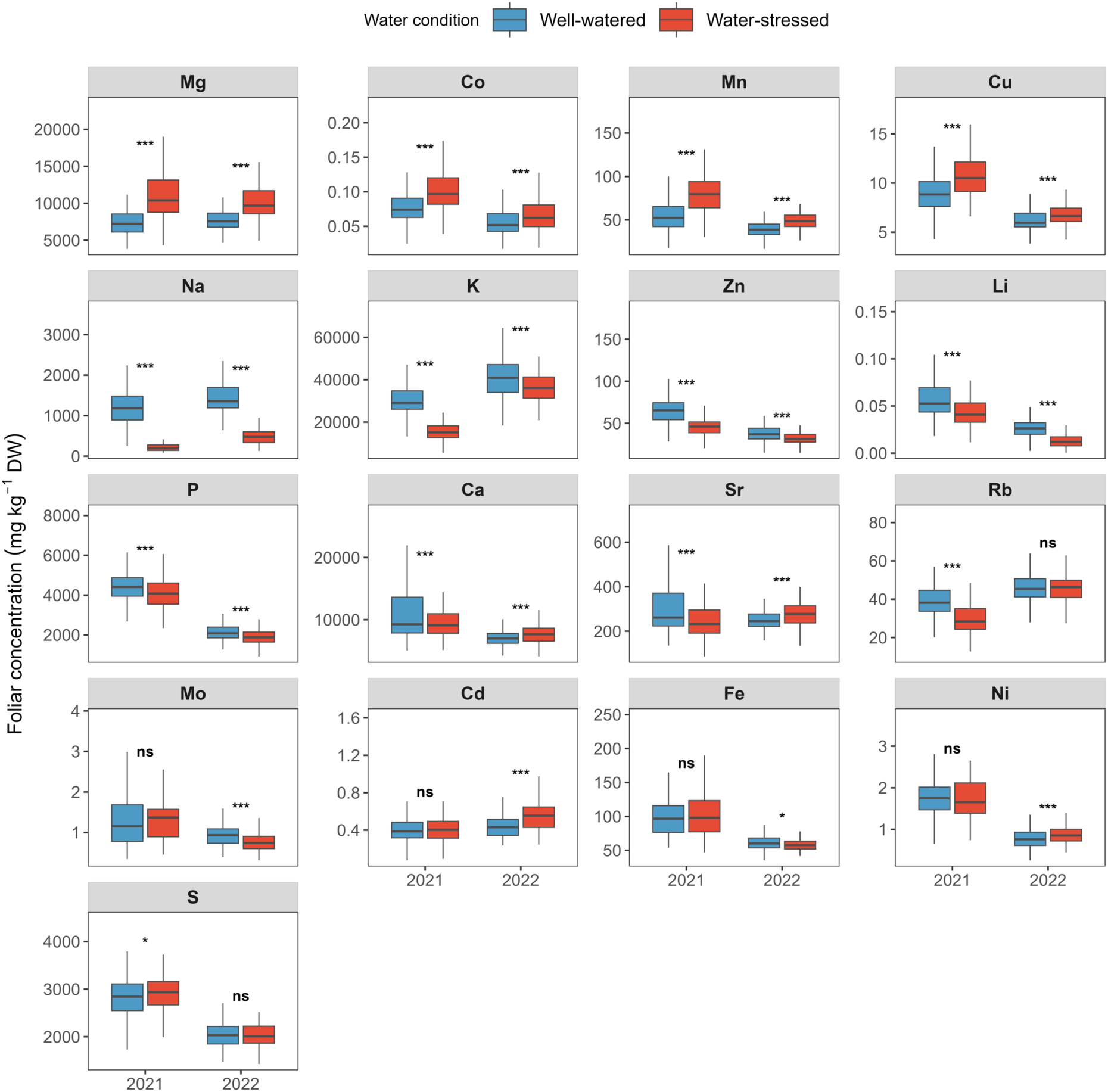
Foliar concentrations of the 17 mineral elements under well-watered and water-stressed conditions, in 2021 and 2022. Boxplots show genotype BLUEs. Asterisks indicate a significant difference between water regimes within a year (paired t-test or Wilcoxon signed-rank test, depending on residual normality; Benjamini-Hochberg FDR-adjusted p-values, corrected within each year: * p<0.05, ** p<0.01, *** p<0.001; ns: not significant).

### Structure of the foliar ionome network across water regimes

To investigate the functional interdependencies underlying foliar ionomic variation, pairwise correlation matrices among the 17 elements were computed separately for each water regime and year (Supplementary Fig. S1). Overall, correlation patterns were largely conserved between well-watered and water-deficit conditions, as indicated by significant Mantel correlations between the two matrices in both years (*r* = 0.88 in 2021; *r* = 0.81 in 2022; *p* = 0.001). However, the proportion of significant pairwise correlations decreased under water deficit, from 66.9% to 59.6% in 2021 and from 44.1% to 25.7% in 2022, indicating an overall weakening of ion-ion associations under stress. Hierarchical clustering on the correlation matrix averaged across all four environments identified two anti-correlated modules: Module 1, comprising thirteen elements (Ca, Cd, Co, Cu, Fe, Li, Mn, Mo, Ni, P, S, Sr, and Zn), and Module 2, comprising four elements (K, Mg, Na, and Rb). This two-module partition was robust to FDR correction at 5% of the correlation matrix. Within Module 1, correlations were observed within the four environments for Ca-Sr (r = 0.86-0.98), Cu-Fe (*r* = 0.66-0.84), Cu-S (*r* = 0.55-0.73), and Cu-P (*r* = 0.56-0.72). Within Module 2, K-Rb was the only pair significant in all four environments (*r* = 0.50-0.66), indicating that Module 2 cohesion is primarily carried by these two elements. Between the two modules, negative correlations were observed as significant in all four environments for Fe-K (*r* = −0.43 to −0.26) and Cu-K (*r* = −0.37 to −0.28), while it was significant in three of the four environments for Rb-Sr, Mo-Na, Cu-Rb and Mg-P. These results show that while the foliar ion correlation network remains structured under early drought, its connectivity and strength are partially reconfigured, reflecting a degree of plasticity in the ionomic covariance structure.

### Relationships between foliar ionome and agro-morphological traits across water regimes

We next examined relationships between individual foliar ion concentrations and agro-morphological traits separately under well-watered and water-deficit conditions, retaining only correlations that were significant in both 2021 and 2022 and with a consistent sign (Supplementary Fig. S2, S3). These correlations were structured according to the two ionomic modules previously identified. Under well-watered conditions, Module 1 elements, particularly Fe, Cu and Mo, were predominantly negatively correlated with phenological traits and plant height, whereas Module 2 elements, particularly K and Na, showed positive correlations with time to flowering and maturity. Under water deficit, the overall contrast between the elements from the two modules was maintained. Cu and Mo were negatively associated with flowering and maturity time, while Fe, Cu, and Mo were negatively associated with plant height. In contrast, K and Mg showed positive associations with flowering and maturity time, and Rb was positively associated with plant height. Relationships with biomass and yield were element- and water-regime-specific, including a positive correlation between Mn and grain weight under well-watered conditions, and negative correlations of Cu and P with shoot dry weight at maturity under water deficit.

We then asked whether, in each of the four environments, correlations observed for individual elements aligned with those observed at the module-level. Composite module scores largely reproduced the contrast already described and extended it to additional traits (Supplementary Table S4). Under well-watered conditions, Module 2 was positively associated with days to flowering, days to maturity, plant height and negatively correlated with harvest index, whereas Module 1 associated negatively with flowering, maturity and plant height, and positively with spike number. Under water deficit, the same contrast persisted for flowering, maturity and plant height, and Module 2 was additionally associated with shoot dry weight at maturity (Supplementary Table S4). The same analysis performed on the combined four environments largely confirmed the main associations identified in the year-specific analyses and revealed additional relationships with grain weight and root anatomical traits under water deficit (Supplementary Tables S5 and S6). The low frequency of significant year interactions indicated that most ion-trait associations were consistent across years.

We next examined whether ion-trait relationships were associated with ionic stability, measured as the stress tolerance index (STI), or with the magnitude of the ionic response to water deficit, measured as the log-ratio index (LR; Supplementary Fig. S4, S5, S6 and S7). The STI-STI correlations recapitulated the major patterns observed for raw ion concentrations, with STI of Fe, Cu and Mo negatively correlated with STI of flowering and maturity time, whereas STI of K showed positive correlations with the STI of the same traits. This phenological pattern was also observed at the module level, with Module 1 negatively and Module 2 positively associated with STI of flowering and maturity time respectively (Supplementary Table S7). LR of Ca and Sr was positively correlated with STI of flowering time and maturity time. No LR-based association was detected at the module level (Supplementary Table S7). Thus, the reproducible ion-trait network was dominated by correlations between ionic stability and phenological stability, while ionic plasticity contributed a smaller, element-specific, set of correlations involving Ca and Sr. A complementary analysis pooling both years, at both the individual-ion and module levels, recovered these correlations and additionally identified relationships involving trait plasticity, not detected under the stricter two-year criterion (Supplementary Table S8). Year interactions were rare among these correlations.

Overall, foliar ion concentrations were consistently associated with flowering and maturity time and plant height, with opposite patterns between the two ionomic modules across years and water regimes. The stability of ion concentrations across water regimes, more than their plasticity, was associated with the stability of phenological traits, plant height and, to a lesser extent, biomass and yield components.

### Multivariate analysis of foliar ionomic variation

Principal component analysis (PCA) of foliar ion concentrations across the four environments resolved two major axes of ionomic variation, together explaining 60.8% of the total variance (PC1, 46.2%; PC2, 14.6%; Fig. 2a). A joint model partitioning each axis by genotype, year, treatment and their interaction showed that PC1 was predominantly associated with year, which accounted for 70.5% of the variance in PC1 scores (*p* < 0.001; Fig. 2b). In contrast, PC2 was predominantly associated with water treatment, which accounted for 65.9 % of the variance in PC2 scores (*p* < 0.001). The main effect of year was not significant for PC2 (*p* = 0.3), suggesting that this axis is predominantly associated with water treatment across years. PC2 was therefore retained as the stress-response axis. The loading structure of PC2 further revealed a coordinated contrast among multiple elements rather than the contribution of a single ion (Na showed the largest positive loading while Mg showed the largest negative loading; Fig. 2c).

**Fig. 2.**
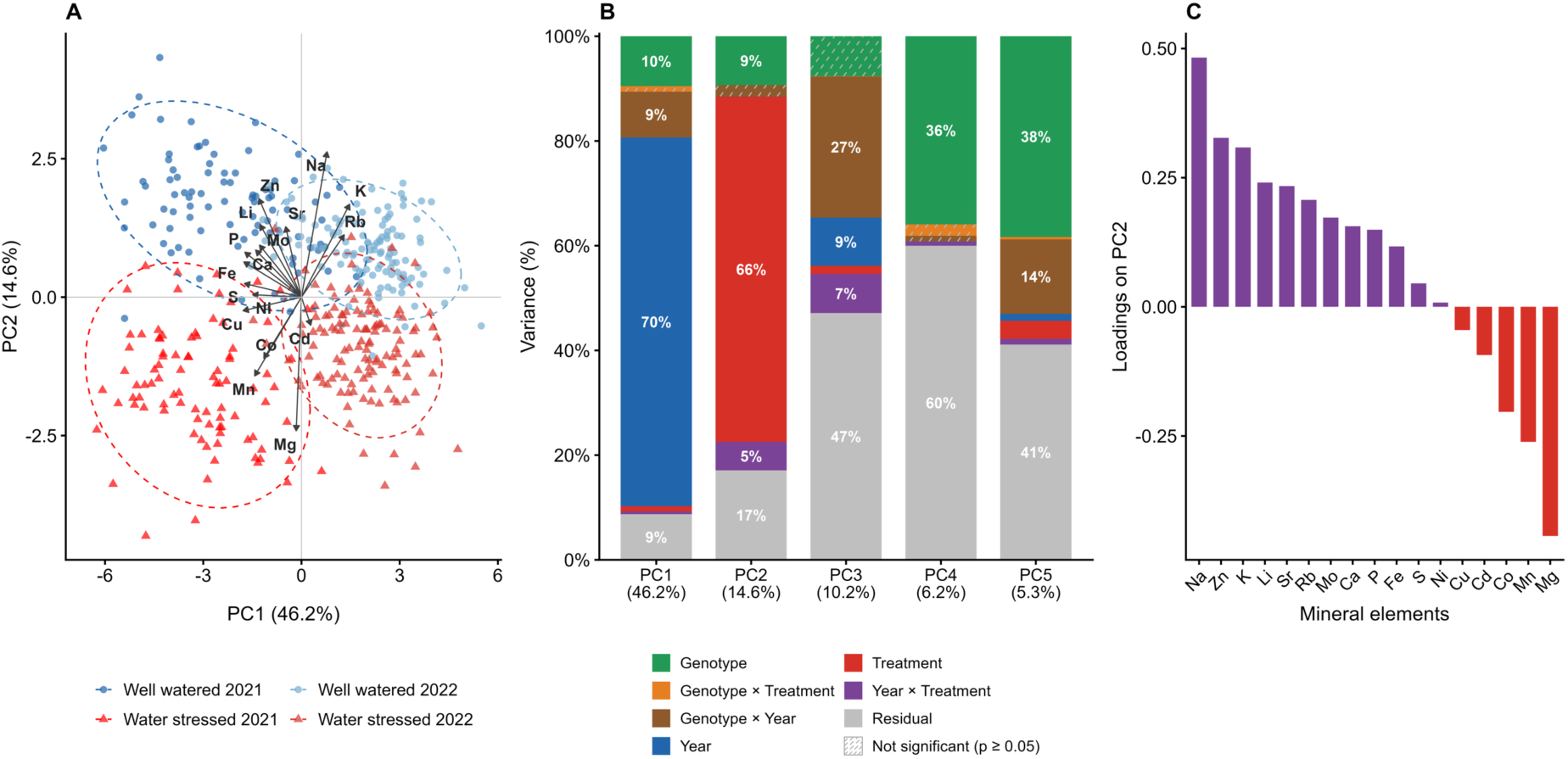
Multivariate structure of the pearl millet leaf ionome under water deficit. (A) Biplot of the first two principal components (PC1, 46.2%; PC2, 14.6%) of the global PCA computed on foliar concentrations (BLUEs) of 17 elements across four environments (well-watered, WW, and water-stressed, WS, in 2021 and 2022). Points are genotype × environment observations, dashed ellipses are 95% ellipses for each environment, and arrows show the element loadings (scaled for display). (B) Percentage of the variance of PC1 to PC5 explained by year, treatment, year × treatment, genotype, genotype × year, genotype × treatment and residual, from sequential sums of squares of a linear model. Hatched segments are not significant (p ≥ 0.05) and percentages are shown for significant terms of at least 4.5%. The percentage of total variance explained by each component is given in brackets. (C) Loadings (eigenvector coefficients) of the 17 elements on PC2, ranked from highest to lowest. Positive loadings (purple) indicate elements higher under well-watered conditions and negative loadings (red) elements higher under water deficit

PC2 also retained a significant genotype component, with genotype accounting for 9.3% of its variance (*p* < 0.001), indicating that genotypes differed significantly in their position along the multivariate ionomic axis associated with water treatment. A significant year × treatment interaction was also detected for PC2 (5.4%, *p* < 0.001), indicating that the magnitude of the ionomic response to water deficit varied between years. No principal component showed a significant G×T variance (*p* ≥ 0.30), whereas G×Y was significant for PC1, PC3 and PC5 (p < 0.01). When PCA variance was partitioned separately by year PC2 remained the component most strongly associated with water treatment in both years, with treatment accounting for 77% of the variance in 2021 and 67% in 2022 (Supplementary Fig. S8).

We next tested whether the ionomic signature associated to response to water stress (PC2) was correlated to the changes in agromorphological traits (LR and STI). The well-watered score (PC2_WW) was negatively correlated with STI for grain weight, plant height and spike weight, while the water-stressed score (PC2_WS) was negatively correlated with STI for spike weight (Supplementary Fig. S9). A model with both years, with year as a fixed covariate and standard errors adjusted for repeated measurement of genotypes, confirmed these five associations and identified new ones (Supplementary Table S9). PC2_WW was additionally negatively correlated with STI for shoot dry weight both at 42/49 DAS and at maturity, and days to flowering and maturity, while PC2_WS was negatively associated with STI for grain weight and tiller number. Associations between PC2 or the stress-induced displacement of the axis (ΔPC2), and trait stability or plasticity were rare.

Altogether, water availability shaped foliar ionomic variation along a single coordinated gradient (PC2). The position of genotypes along this gradient under well-watered conditions, more than the shift induced by water deficit, was associated with STI for shoot biomass, grain weight, spike weight and plant height, suggesting that the leaf mineral profile of unstressed plants could help identify tolerant genotypes.

### Ionomic prediction of genotype-level drought tolerance

Having established that the PC2 axis under well-watered conditions was associated with STI for biomass and grain weight, we tested whether the ionome measured under well-watered conditions could predict drought tolerance in pearl millet. Three predictors were compared: (i) the well-watered score of PC2 (PC2_WW), (ii) the single ion most strongly correlated with the trait, and (iii) a multi-ion signature obtained by penalized regression.

For STI grain weight, PC2_WW showed the highest accuracy, ahead of the multi-ion signature and the single ion selected within training sets (mean cross-validated r = 0.300, 0.197 and 0.195, respectively), suggesting a better prediction of grain weight maintenance (Supplementary Table S10). For STI shoot dry weight at maturity, the multi-ion signature, which retained Mg, Fe and Mo in all 20 refits, significantly outperformed the single ion and PC2_WW (r = 0.422, 0356 and 0.317, respectively), indicating improved prediction from a specific multi-ion signature (Supplementary Table S10).

The tercile analysis identified 42 observed tolerant and 42 observed sensitive genotypes whose predicted scores differed significantly for both traits (Wilcoxon, p = 0.015 and p = 1.7×10⁻¹¹, respectively; Supplementary Table S10). Classification in each group for shoot biomass and grain weight was non-random (Fisher’s exact test, OR = 8.2, p = 3×10⁻⁷), genotype ranking was conserved (Spearman ρ = 0.54, *p* < 2.2×10⁻¹⁶), and similar for observed STI (OR = 4.2, p = 5×10⁻⁴; ρ = 0.38, *p* = 1×10⁻⁵), with 27 genotypes jointly classified as tolerant for shoot biomass and grain weight (Supplementary Table S11). Predicted scores were not substantially confounded by population structure, which was limited in this panel (genomic PC1 explained 4.6% of the variance).

Overall, foliar ionomic profiles measured on well-watered plants predicted the maintenance of grain weight and biomass under water deficit. The most informative predictor differed between traits, the global PC2 axis for grain weight and a multi-ion model consistently retaining Mg, Fe and Mo across refits for biomass. Based on predicted scores, 27 genotypes were classified as tolerant for both traits, identifying drought-tolerant germplasm and illustrating the use of pre-stress ionomic profiling for early, non-destructive screening.

### Genetic variance in foliar ion accumulation and its interaction with water treatment

To quantify the respective contributions of genetic and environmental factors to foliar ion accumulation, linear mixed-effects models were fitted for each element, with water regime, year, and their interaction as fixed effects, and genotype, genotype by treatment (G×T), and genotype by year (G×Y) as random effects. The random genotype effect was significant for multiple ions, reflecting substantial genetic variation in mean foliar concentrations independently of water regime. The proportion of total phenotypic variance attributable to genetic effects ranged widely across elements (from 21% for Zn to 40.6% for Cu; Fig. 3; Supplementary Table S12). Among all elements analysed, only potassium showed a significant G×T interaction (*p* = 5.48×10⁻³), accounting for 15.9% of its total variance (Supplementary Table S12), indicating that pearl millet genotypes differ specifically in their foliar potassium content in response to early water deficit. Drought-induced changes in the other elements analysed were largely independent of the genotypes.

**Fig. 3.**
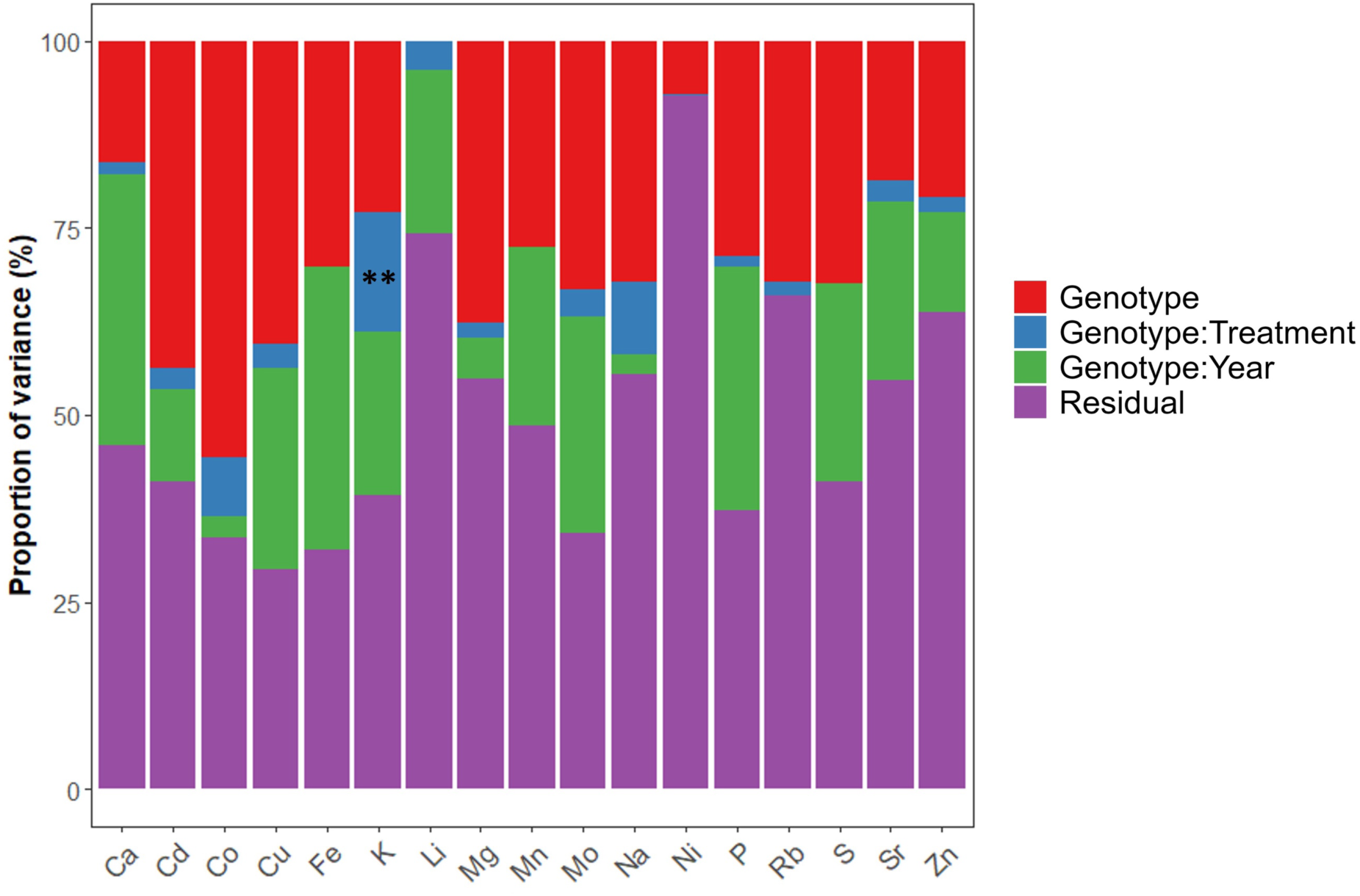
Contribution of variance components to total variance of leaf ion concentrations. Variance is partitioned among random effects (genotype, treatment, genotype × treatment interaction), expressed as a percentage of total trait variance. Asterisks indicate significant genotype × treatment interactions (*p* < 0.05).

### Constitutive genetic architecture of the pearl millet foliar ionome

To identify markers with effects consistent across all four environments, genome-wide association analysis was performed using the metaGE meta-analysis framework without contrast for treatments. Significant loci were identified for 12 of the 17 quantified elements, yielding 48 significant SNPs that resolved into 42 independent QTLs after grouping markers within linkage disequilibrium blocks (Table 2; Supplementary Table S13). Four instances of co-localisation between markers associated with different elements were observed. The SNP CM007983_27393436 on chromosome 2 was associated with both K and Cu accumulation, the SNP CM007986_6220019 on chromosome 5 was associated with both Fe and Ca accumulation, and the SNP LKME02014639.1_317356 on an unanchored contig and CM007983_157814421 on chromosome 2 were associated with both K and Rb accumulation (Table 2; Supplementary Table S13). Among all elements, K showed the broadest chromosomal distribution of associated loci. Genes present within the linkage disequilibrium (LD) block surrounding each QTL and the Z-scores of all significant SNPs in each environment are summarised in Supplementary Table S13. These 42 QTLs thus represent loci with consistent effects on the foliar ionome across all four year × water regime combinations (well-watered and drought-stressed).

**Table 2.** Constitutive QTLs for leaf element concentrations detected by multi-environment GWAS meta-analysis across four environments in the PMiGAP pearl millet panel.

| **Element** | **Markers** | **QTLs** | **Chromosomes** | **Co-localised QTLs** |
|---|---|---|---|---|
| Ca | 4 | 4 | 1, 3, 5, contig | 1 (Fe) |
| Cd | 2 | 1 | 1 | – |
| Cu | 5 | 3 | 2, 3, 5 | 1 (K) |
| Fe | 5 | 4 | 2, 5, 7 | 1 (Ca) |
| K | 9 | 8 | 1, 2, 6, 7, contig | 3 (Cu, Rb) |
| Li | 5 | 4 | 2, 4, contig | – |
| Mg | 2 | 2 | 3, contig | – |
| Mn | 2 | 2 | 2, contig | – |
| Mo | 1 | 1 | 6 | – |
| Na | 2 | 2 | 3, 4 | – |
| Rb | 10 | 10 | 1, 2, 6, contig | 2 (K) |
| Zn | 1 | 1 | 2 | – |

Markers: total number of significant SNPs (*p* < 10<sup>−5</sup>) from the metaGE fixed-effect meta-analysis without contrast, performed for four GWAS methods (LFMM2, EMMA, BLINK and CMLM). QTLs: number of QTL associated; markers of the same element located within the same LD region (r² > 0.5) were grouped into one QTL. Chromosomes: chromosomal positions of the identified QTLs; contig refers to unanchored genomic contigs that could not be assigned to a specific chromosome. Co-localised QTLs: number of QTLs sharing a marker with a QTL for another element (element in brackets). Details for each marker are given in Supplementary Table S13.

Markers: total number of significant SNPs (*p* < 10⁻⁵) from the metaGE fixed-effect meta-analysis without contrast, performed for four GWAS methods (LFMM2, EMMA, BLINK and CMLM). QTLs: number of QTL associated; markers of the same element located within the same LD region (r² > 0.5) were grouped into one QTL. Chromosomes: chromosomal positions of the identified QTLs; contig refers to unanchored genomic contigs that could not be assigned to a specific chromosome. Co-localised QTLs: number of QTLs sharing a marker with a QTL for another element (element in brackets). Details for each marker are given in Supplementary Table S13.

### Identification of QTLs controlling leaf potassium plasticity

We next looked for QTL controlling ionomics plasticity in response to vegetative water stress. Among the 17 elements, K was the only one with a significant genotype × treatment variance component (*p* = 5.48×10⁻³). To dissect the genetic architecture of potassium plasticity in response to water deficit, three complementary analyses were applied: (i) condition-specific GWAS within each water regime, (ii) a genotype by treatment contrast model in MetaGE within all four environments, and (iii) GWAS on the potassium plasticity index LR_K. These analyses identified a set of loci whose effects are either condition specific or relate to the potassium response to water stress.

Under well-watered conditions, five significant SNPs were detected on chromosomes 2, 3, 6 and 7, resolving into five independent QTLs (Supplementary Table S14). Among these, CM007983_27393436 on chromosome 2 (qKC-2a) and CM007987_104928234 on chromosome 6 (qKC-6b) were also identified in the four-environment analysis without contrast for treatments. Under water stress, two significant SNPs were detected: CM007987_17176446 on chromosome 6 (qKC-6a) and LKME02014639.1_317356 on an unanchored contig (qKC-uc1). Both SNPs were also identified in the four-environment analysis without contrast, with Z-scores of greater magnitudes under water stress than under well-watered conditions. Three SNPs detected under well-watered conditions were absent from both the four-environment and water stress analyses and defined well-watered-specific QTLs: CM007984_207071465 on chromosome 3 (qKE-3), CM007988_37716247 on chromosome 7 (qKE-7a) and CM007988_136644636 on chromosome 7 (qKE-7b).

In the genotype by treatment contrast model applied across four environments, a single SNP reached significance: CM007983_27393436 on chromosome 2 (qKC-2a), already identified in the four-environment without contrast analysis, and well-watered analyses (Supplementary Table S14). This SNP had Z-scores of the same sign in all four environments, but larger under well-watered than under water stress conditions (Z = 5.34 and 4.19 under well-watered conditions; Z = 3.29 and 2.76 under water stress, for 2021 and 2022 respectively). The C allele was associated with higher K concentration under both water regimes, with a larger difference under well-watered conditions (C/C vs T/T: 39,689 vs 32,294 mg kg⁻¹) than under water stress (28,114 vs 24,322 mg kg⁻¹). The SNP class × treatment interaction on K concentration was significant (*p* = 0.002), i.e. the effect of qKC-2a differed in size between water regimes but not in direction.

Direct GWAS on potassium plasticity index LR_K identified two significant SNPs on chromosome 6: CM007987_110250858 and CM007987_110250945, located 87 bp apart within the same linkage disequilibrium block (Supplementary Table S14). Together they define a single QTL designated qKP-6. Neither SNP was detected in previous GWAS on absolute potassium concentrations. Pairwise linkage disequilibrium between CM007987_110250858 and all other significant potassium SNPs was r² < 0.09 regardless of chromosomal position, indicating that qKP-6 is distinct from all previously identified potassium loci. G/G genotypes at the CM007987_110250858 locus accumulated more potassium than A/A genotypes under well-watered conditions, whereas potassium concentrations did not differ significantly between genotypes under water stress. A/A genotypes showed a mean potassium loss of 27.4% under water stress relative to well-watered conditions, compared to 40.1% for G/G genotypes. A significant SNP class × treatment interaction (*p* = 2.4×10⁻⁴) indicated that allelic effects at this locus depend on the water regime. Because LR_K was associated with flowering time (*p* = 1.6×10⁻⁴) and A/A genotypes flowered 4.4 days earlier than G/G genotypes (*p* = 0.003), models were additionally adjusted for flowering time. The qKP-6 effect on LR_K remained significant (*p* < 0.001; ΔR²m = 0.037), with a 14.7% reduction between homozygous classes, whereas the effect on potassium concentration under well-watered conditions was no longer significant (*p* = 0.10), suggesting that the association with K plasticity was not fully accounted for by flowering time. The effect on LR_K was significant in each year (2021: *p* = 0.017; 2022: *p* = 6.5×10⁻⁵). Altogether, our analyses identified a new QTL, qKP-6, associated with the plasticity of leaf K concentration under vegetative drought stress and distinct from the loci for K concentration.

### Identification of a candidate gene in qKP-6

At qKP-6, linkage desequilibrium analysis around SNP CM007987_110250858 identified a 14.3-kb LD block spanning 110,236,620 to 110,250,945 bp and containing CM007987_110250945, a second SNP significantly associated with the potassium plasticity index (Figure 4; Supplementary Table S14). The block also encompassed several additional variants, including four SNPs located within *Pgl_GLEAN_10012096* (110,235,838-110,237,176 bp) which partially overlaps the LD block. These four SNPs showed strong LD with the index SNP (*r²* = 0.77-0.86), supporting *Pgl_GLEAN_10012096* as the strongest positional candidate at this locus. BLASTp analysis of the predicted 348-amino-acid protein revealed sequence similarity to uncharacterised proteins from related grass species, including *Urochloa humidicola* (59% sequence identity), *Setaria italica* (52%) and *Paspalum vaginatum* (46%). InterProScan identified no recognisable functional domains in the predicted protein. Thus, *Pgl_GLEAN_10012096* is prioritised as a positional candidate for potassium plasticity at qKP-6, although its molecular function remains unknown.

**Fig. 4.**
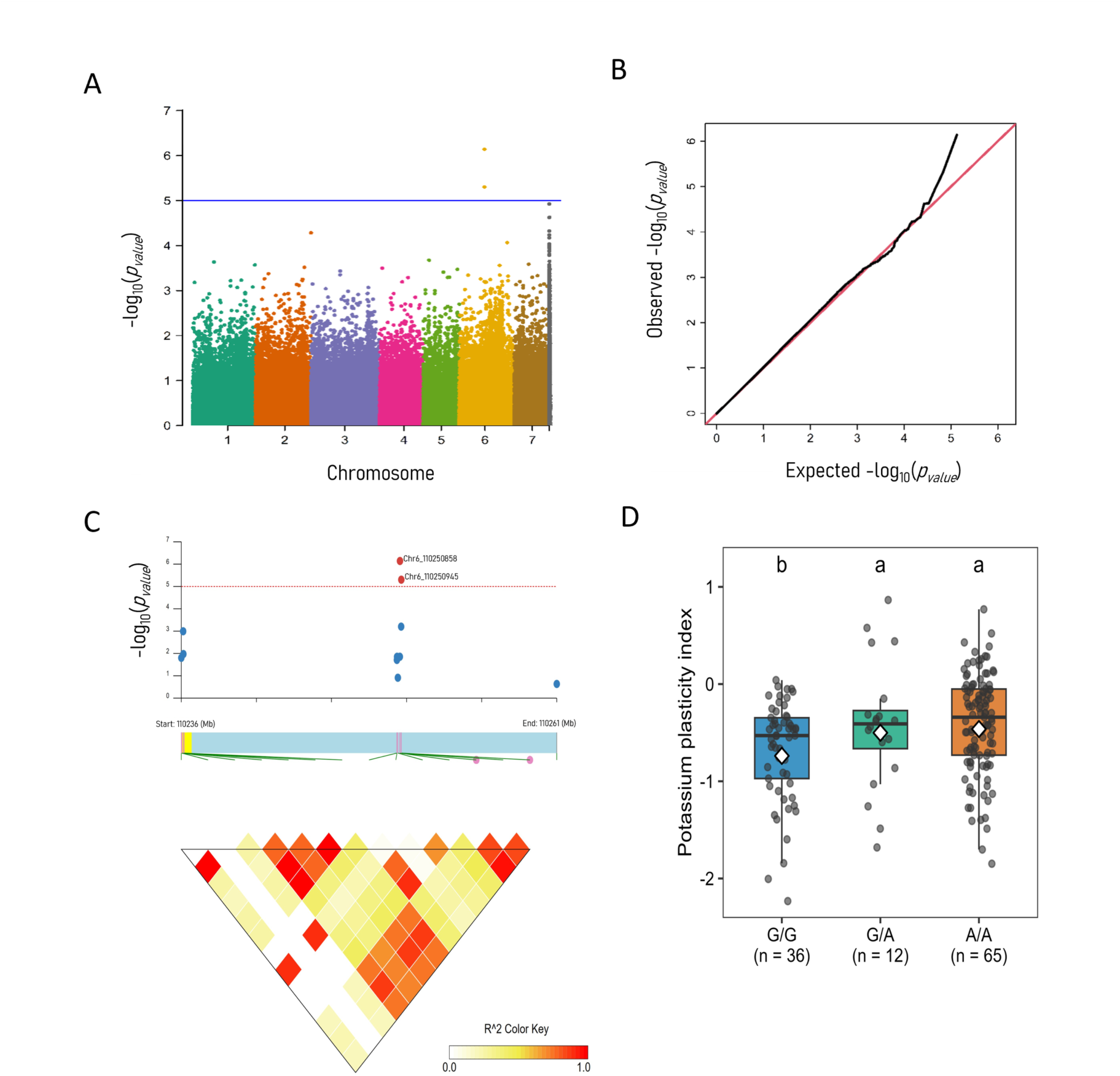
Genome-wide association analysis of foliar potassium plasticity and characterization of the qKP-6 locus on chromosome 6. (A) Manhattan plot of -log₁₀(p) values for all SNPs from the GWAS of the potassium plasticity index (LR_K). Colours distinguish chromosomes 1 to 7 and unanchored contigs (grey). The blue line indicates the genome-wide significance threshold (*-log₁₀ p = 5*). (B) Quantile-quantile plot of observed against expected -log₁₀(p) values. The red line is the expectation under the null hypothesis. (C) Regional Manhattan plot of the significant locus on chromosome 6 (110,236 to 110,261 kb), with the two significant SNPs in red and the local linkage disequilibrium (LD) structure. Pairwise LD (r²) values among SNPs are displayed as a colour gradient from 0 to 1. The yellow segment marks the transcribed region of *Pgl_GLEAN_10012096.* (D) LR_K by allelic class at CM007987_110250858 (G/G, G/A, A/A; n = number of lines). Points are line × year observations, boxes show the median and interquartile range, and white diamonds the means. Different letters indicate significant differences between classes (*p <* 0.05; Tukey-adjusted pairwise contrasts of estimated marginal means from the linear mixed model with year as fixed effect and genotype as random effect).

## Discussion

In this study, we investigated the drought response of the leaf ionome in pearl millet, aiming to identify ions associated with early drought tolerance and to evaluate whether plasticity in ion content contributes to this tolerance. In addition, we searched for genetic determinants associated with ion content stability across water regimes and with drought-induced plasticity, with the aim of supporting breeding towards improving pearl millet resilience to drought in terms of nutrient use efficiency and forage quality. Our findings suggest that the stability of several leaf ions is associated with drought tolerance in pearl millet. We further identified the genetic architecture underlying both ion stability across water regimes and drought-induced ion plasticity highlighting potassium as a particularly variable component of the drought response.

Leaf ionomic composition has been shown to be remodelled by drought stress in several crops (Fischer *et al*., 2019; D’Oria *et al*., 2022). In young wheat leaf blades, moderate to severe drought stress induced a significant reduction in the concentration of most ions (D’Oria *et al*., 2022), whereas in maize cobs, mild drought increased ion concentrations, while severe drought caused a strong reduction (Fischer *et al*., 2019). Substantial variation among genotypes has also been observed in ionomic responses to drought (D’Oria *et al*., 2021, 2022; Quagliata *et al*., 2025). In our study, the use of a large panel of pearl millet genotypes confirmed that the leaf ion content under drought varied largely among genotypes and that this variation was partly under genetic control. The strongest source of environmental variation in ionomic composition was associated with the year, as observed in the principal component analysis. The effect of the drought treatment was secondary and was mainly captured by the second principal component axis. Despite the strong year effect, correlations among ions observed under irrigated conditions were generally conserved under drought stress, although they tended to weaken. Indeed, two covariation modules were consistently identified across conditions: one comprising Ca, Cd, Co, Cu, Fe, Li, Mn, Mo, Ni, P, S, Sr and Zn, and another comprising Na, K, Rb and Mg. The concentrations of some of these ions were consistently affected by drought across both years: Na, K, Zn and Li concentrations decreased, whereas those of Mg, Co, Mn and Cu increased. Together, these results suggest that water deficit does not disrupt the overall structure of the leaf ionome but rather alters elemental concentrations while largely preserving their covariation structure. This interpretation is consistent with the concept of ionomic homeostasis, whereby coordinated relationships among elements can be maintained despite changes in their absolute concentrations under water deficit (Salt *et al*., 2008).

The concentrations of several ions have been associated with improved drought tolerance in crops. In alfalfa, changes in the concentrations of ions such as Na, K and Ca were associated with the maintenance of cellular homeostasis and water balance under drought (Chen *et al*., 2026). Furthermore, several studies have illustrated the beneficial role of K in mitigating the effects of drought stress in wheat (Alzahrani *et al*., 2026), barley (Feng *et al*., 2020) or poplar (Zhang *et al*., 2023). In line with these findings, our results showed that Na and K concentrations were positively correlated with time to flowering, while parameters describing the ion module containing these two elements were also positively correlated with shoot dry weight at maturity. However, Na and K concentrations were significantly reduced under drought. Such reductions may result from decreased K uptake and translocation to the shoots due to drought-induced limitations in transpiration and may illustrate a potential trade-off between water conservation and nutrient uptake required for osmoregulation (Nieves-Cordones *et al*., 2014, 2019). Conversely, Cu and Fe concentrations were negatively associated with plant height and shoot dry weight at maturity under drought. Similarly, although Mg concentration was not directly correlated with agro-morphological traits, the computation of an ionomic signature based on a principal component analysis revealed that the water stress score (PC2_WS), to which Mg was a major contributor, was negatively associated with the stress tolerance index (STI) for grain weight, tiller number and shoot dry weight at maturity. Leaf Mg and Cu concentrations increased under drought and, although Fe concentration did not increase significantly in these conditions, it was closely associated with Cu concentration. This pattern is consistent with the possibility that a variation in Cu and Fe levels contributes to oxidative stress under drought conditions (Ravet and Pilon, 2013). Therefore, both ions deficiency (likely Na and K) and excessive accumulation of specific ions (Mg, Cu and Fe) may have contributed to the reduction in shoot biomass production and yield under drought. However, these associations should be interpreted as relationships rather than evidence of direct causal effects. Beyond individual ion associations, the constitutive foliar ionome under well-watered conditions also distinguished tolerant and sensitive genotypes. Grain yield maintenance under drought tended to be best captured by the well-watered score (PC2_WW). Biomass tolerance, in contrast, was better predicted by a sparse Mg-Fe-Mo signature than by PC2_WW, indicating that a combination of elements carried more information than any single elemental concentration. The significant, though partial, overlap between genotypes tolerant for both traits is consistent with a shared tolerance component, and the resulting dual-tolerant genotypes provide promising candidates for subsequent field validation.

Phenotypic plasticity can play important roles in the ability of plants to cope with drought stress (Schneider, 2022). However, adaptive responses to drought require appropriate coordination, as maladaptive plasticity may negatively affect plant growth and fitness (Brooker *et al*., 2022). In our study, no correlations were observed between the plasticity of individual ions, or of the leaf ionome, and the plasticity of agro-morphological traits. In contrast, the stability of leaf ion concentration across water regimes was associated with the stability of agronomic traits. These results suggest that, in pearl millet, adaptation to drought may rely more on the maintenance of ion homeostasis than on the adaptive changes in leaf ion content. Correlation analyses further suggest that this strategy may be beneficial under drought for some ions, such as Na and K to avoid a reduction in leaf concentration, but also Mg and Cu to avoid an increase in leaf concentration. Interestingly, among all the ions studied, only leaf K concentration showed a significant genotype by treatment interaction, which explained 15.9 % of its variance. This suggests that the ionomic reorganisation induced by drought may represent a conserved physiological response in pearl millet. In contrast, the genotype-specific response of K indicates that this otherwise conserved response contains a heritable component that may be exploitable for breeding. Genotype-dependent K response may reflect its multifunctional role in plant water relations and growth, including osmotic adjustment, stomatal regulation or transport (Mishra *et al*., 2026). Genetic variation in these processes may therefore contribute to the differences observed among genotypes in K homeostasis under drought stress.

K was identified as being associated with improved drought tolerance, although its concentration tended to decrease under drought. Therefore, QTL contributing to the maintenance of K concentration across water regimes may be particularly relevant for breeding. GWAS identified eight constitutive loci (qKC) for K concentration that were not affected by water treatment. Interestingly, two of these loci (qKC-6a and qKC-uc1) were also identified specifically for K concentration under drought stress, suggesting that they may be of particular interest for increasing K concentration constitutively across water regimes. Because the drought response of leaf K concentration was genetically controlled, we used plasticity indices to quantify this response and applied genetic models based on meta-analysis of contrasts between water treatments to identify its genetic basis. Among the different plasticity indices, we selected the ratio, which has been shown to be suitable for uncovering the genetic architecture of plasticity (Arenas *et al*., 2025). One locus on chromosome 6 (qKP-6) was identified as associated with this index. The alternate allele limits K depletion under drought without affecting constitutive K levels. Although no agro-morphological trait showed a significant association with this SNP that could reach genome-wide significance in GWAS analyses, the favourable allele was significantly associated with greater stability of flowering time under drought. This association raises the possibility that qKP-6 contributes to coordinated regulation of K homeostasis and developmental stability under drought, although the underlying mechanism remains to be established. Within the LD block surrounding qKP-6, the only annotated gene (*Pgl_GLEAN_10012096*) encodes a protein of unknown function. Its strong LD with the lead SNP makes it a positional candidate, but functional validation will be required to determine whether it directly contributes to K plasticity. The identification of these QTL provides promising opportunities for improving drought tolerance through the manipulation of leaf K concentration.

## Conclusion

Pearl millet is a cereal crop of major importance for food security, particularly in the driest regions of the world. Accelerating its genetic improvement is crucial to maintain food security under current and future climate conditions. Our results suggest that the response of pearl millet ionome to drought involves the maintenance of ion homeostasis, while highlighting the central role of K both in plant growth and in drought tolerance. The identification of distinct QTL associated with K accumulation and its plasticity in response to drought provides new insights into the genetic basis of K nutrition and plasticity in pearl millet, opening new prospects for breeding. Ultimately, these findings contribute to a better understanding of the mechanisms underlying plant adaptation to drought, a critical challenge in the context of climate change.

## Acknowledgements

This work was supported by the European Plant Phenotyping Network (EPPN2020 project Ionomil n°386), the Royal Society (Anatomics grant ICA-R1-180356 to MB and NK) and the Agence Nationale de la Recherche (ANR-20-CE20-0016 to AG). P.N.G. was supported by a joint PhD grant from the Institut de Recherche pour le Développement (IRD, France) and the French Embassy in the Central African Republic.

## Data availability

Data are available in the DataSuds database (https://doi.org/10.23708/DWGEAJ).

