## Supplemental figures for "Leaf ionomic stability shapes drought adaptation in pearl millet"

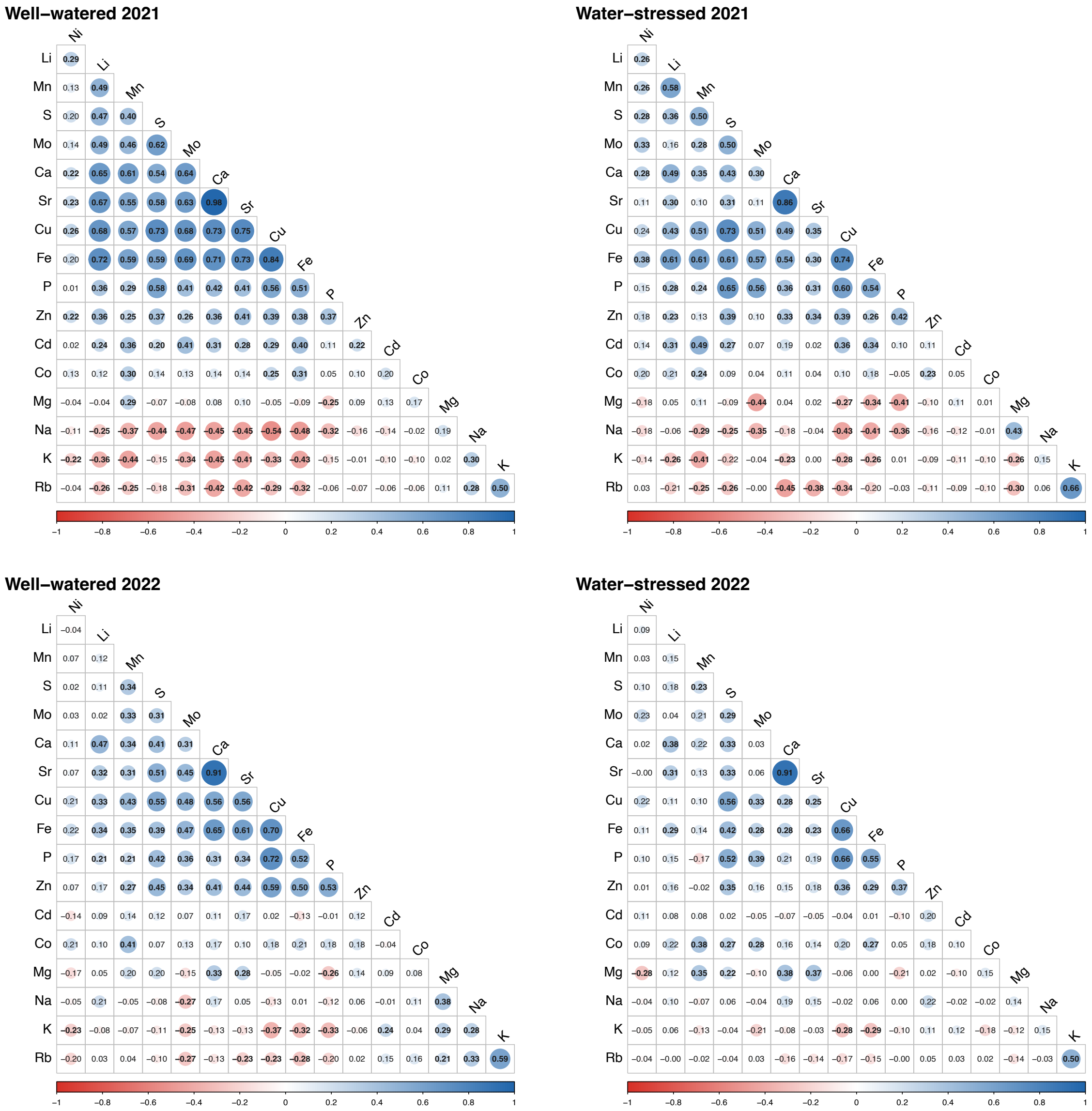


**Fig. S1. Pairwise Pearson correlation matrices among 17 foliar mineral elements, by environment.** Correlations were computed separately for well-watered (WW) and water-deficit (WS) conditions in 2021 and 2022, on raw BLUEs (136 element pairs per environment). Circle size and colour intensity are proportional to the correlation coefficient (r).

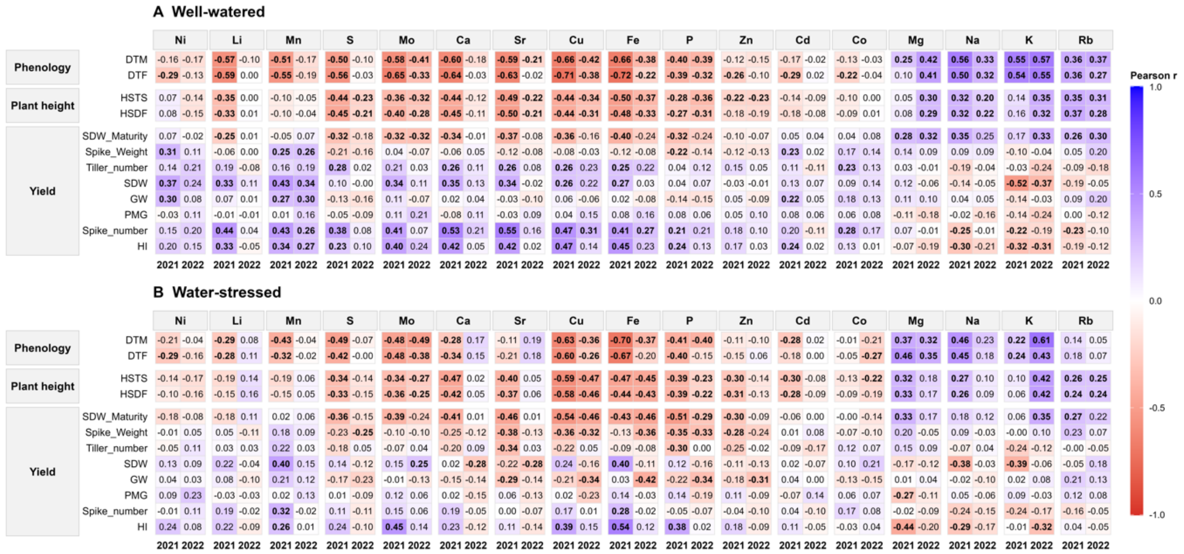


**Fig. S2. Pairwise Pearson correlations between foliar ion concentrations and agromorphological traits (phenology, plant height and yield components), under well-watered (A) and water-stressed (B) conditions.** For each element and trait, correlation coefficients (Pearson r) are shown separately for 2021 and 2022. Values are displayed in all cells; bold values indicate correlations significant after Benjamini-Hochberg FDR correction ($p < 0.05$), applied within each year, water regime and trait category. Colour scale reflects the sign and magnitude of r, from red (negative) to blue (positive). Trait abbreviations are defined in Supplementary Table S1.

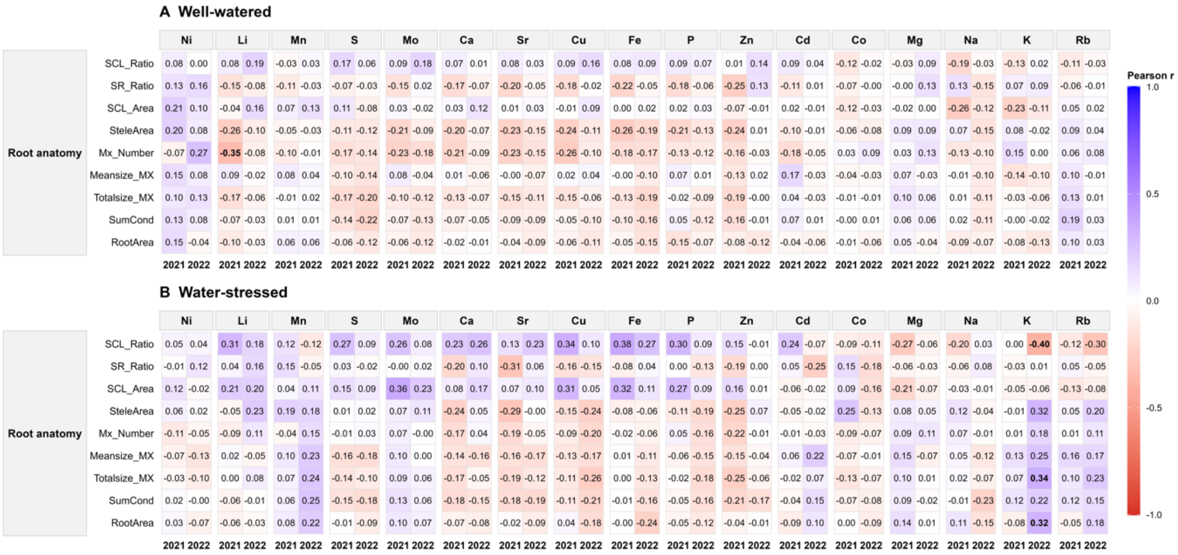


**Fig. S3. Pairwise Pearson correlations between foliar ion concentrations and root anatomical traits, under well-watered (A) and water-stressed (B) conditions.** For each element and trait, correlation coefficients (Pearson r) are shown separately for 2021 and 2022. Values are displayed in all cells; bold values indicate correlations significant after Benjamini-Hochberg FDR correction ($p < 0.05$), applied within each year, water regime and trait category. Colour scale reflects the sign and magnitude of r, from red (negative) to blue (positive). Trait abbreviations are defined in Supplementary Table S1.

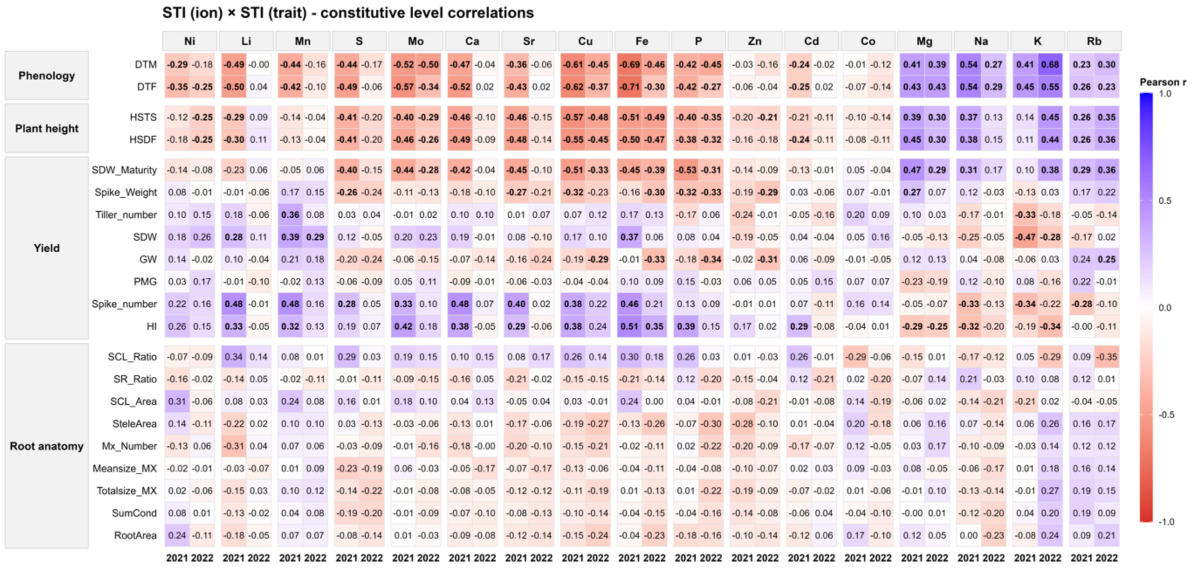


**Fig. S4. Pairwise Pearson correlations between ionic stress tolerance (STI) and the corresponding STI of agromorphological and root traits.** For each element and trait, correlation coefficients (Pearson r) are shown separately for 2021 and 2022, across phenology, plant height, yield components and root anatomical traits. Values are displayed in all cells; bold values indicate correlations significant after Benjamini-Hochberg FDR correction ($p < 0.05$), applied within each year, index combination and trait category. Colour scale reflects the sign and magnitude of r, from red (negative) to blue (positive). Trait abbreviations are defined in Supplementary Table S1.

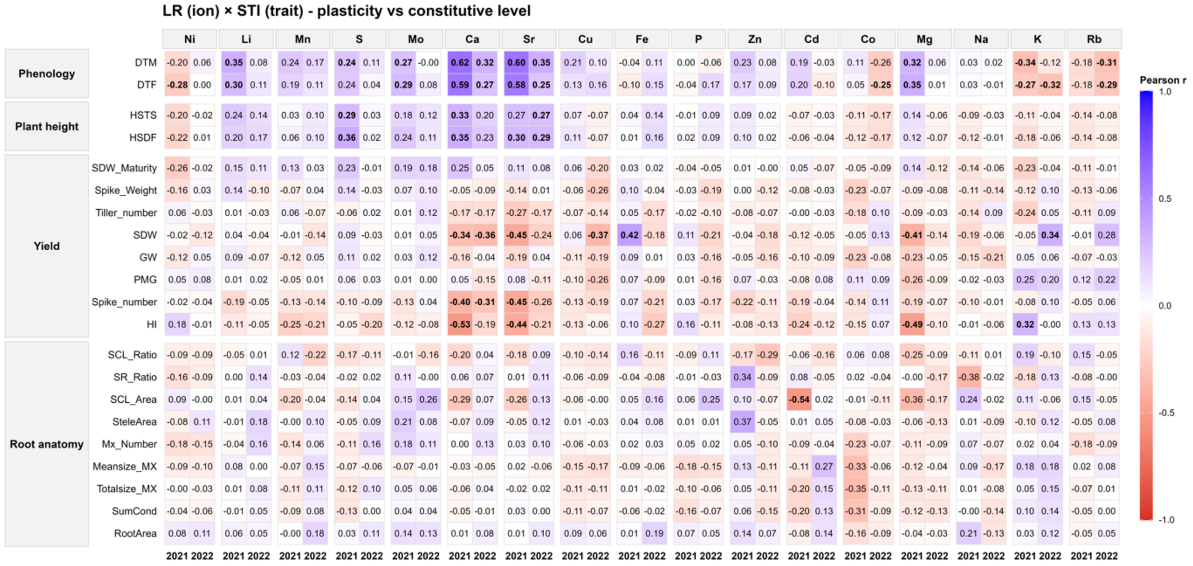


**Fig. S5. Pairwise Pearson correlations between ionic plasticity (LR) indices and the corresponding STI of agromorphological and root traits.** For each element and trait, correlation coefficients (Pearson r) are shown separately for 2021 and 2022, across phenology, plant height, yield components and root anatomical traits. Values are displayed in all cells; bold values indicate correlations significant after Benjamini-Hochberg FDR correction ($p < 0.05$), applied within each year, index combination and trait category. Colour scale reflects the sign and magnitude of r, from red (negative) to blue (positive). Trait abbreviations are defined in Supplementary Table S1.

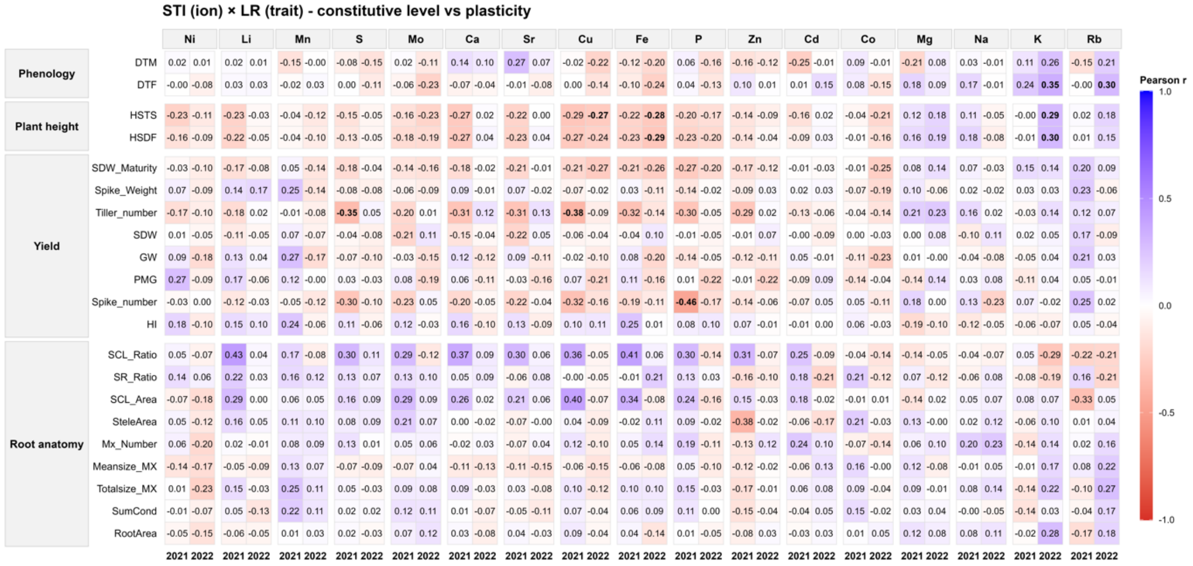


**Fig. S6. Pairwise Pearson correlations between ionic stress tolerance (STI) and the corresponding plasticity (LR) indices of agromorphological and root traits.** For each element and trait, correlation coefficients (Pearson r) are shown separately for 2021 and 2022, across phenology, plant height, yield components and root anatomical traits. Values are displayed in all cells; bold values indicate correlations significant after Benjamini-Hochberg FDR correction ($p < 0.05$), applied within each year, index combination and trait category. Colour scale reflects the sign and magnitude of r, from red (negative) to blue (positive). Trait abbreviations are defined in Supplementary Table S1.

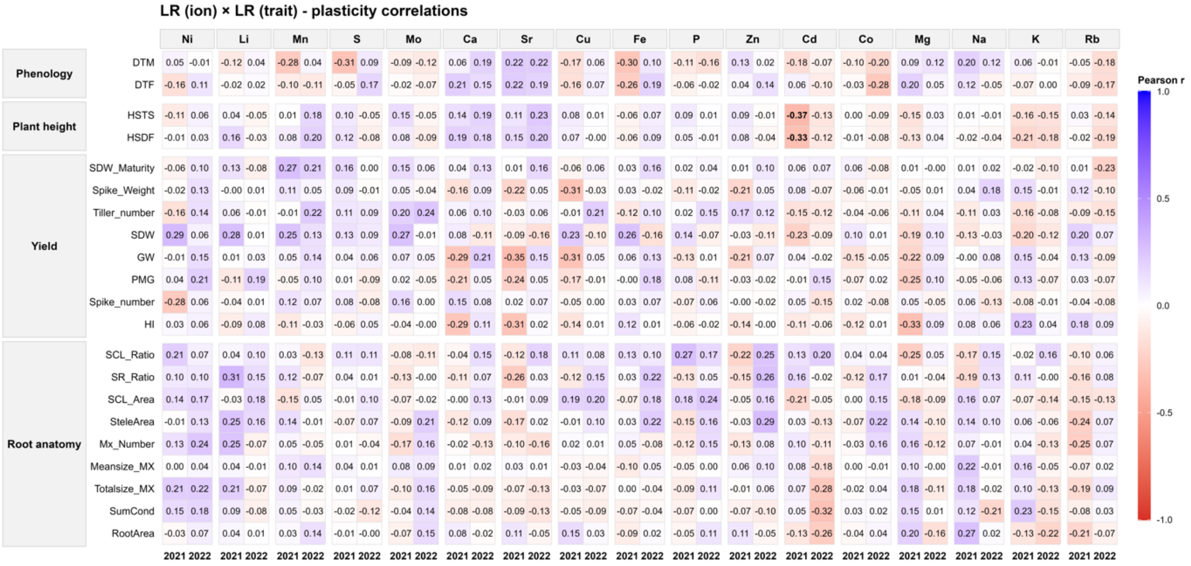


**Fig. S7. Pairwise Pearson correlations between ionic plasticity (LR) indices and the corresponding LR indices of agromorphological and root traits.** For each element and trait, correlation coefficients (Pearson r) are shown separately for 2021 and 2022, across phenology, plant height, yield components and root anatomical traits. Values are displayed in all cells; bold values indicate correlations significant after Benjamini-Hochberg FDR correction ($p < 0.05$), applied within each year, index combination and trait category. Colour scale reflects the sign and magnitude of r, from red (negative) to blue (positive). Trait abbreviations are defined in Supplementary Table S1.

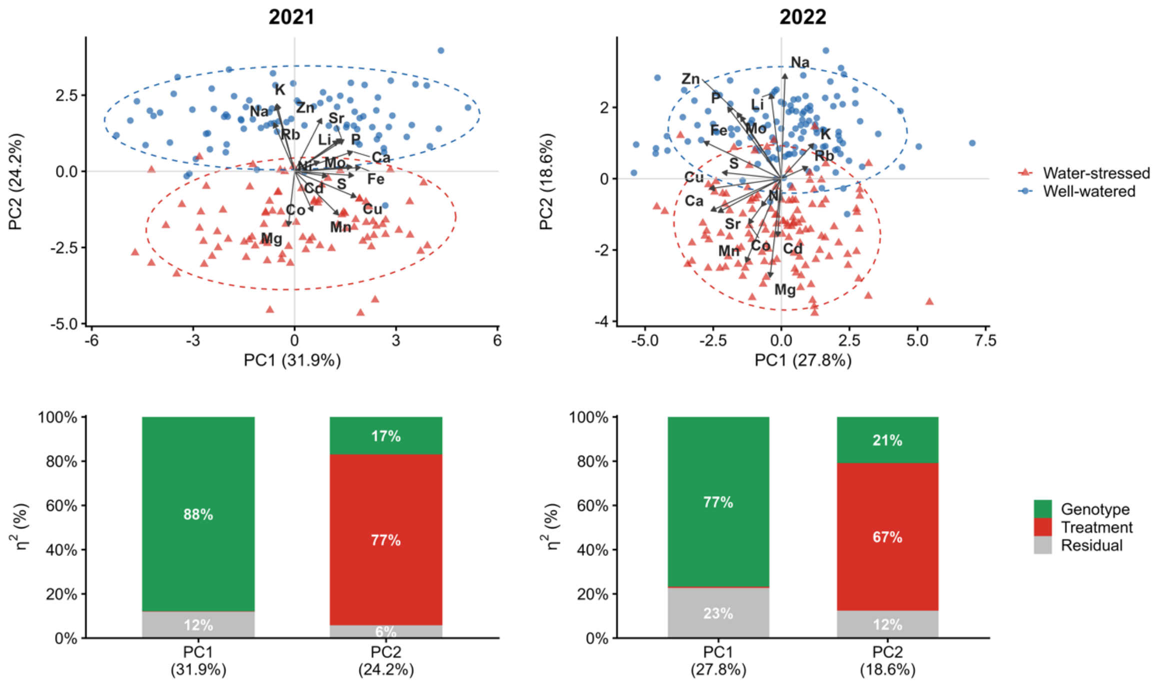


**Fig. S8. Year-specific principal component analysis of the foliar ionome**. Upper panels, biplots of the first two principal components of the PCA computed separately for 2021 (left) and 2022 (right) on the 17 elements measured under well-watered (blue circles) and water-stressed (red triangles) conditions. Dashed ellipses enclose 95% of the observations of each treatment and arrows show the element loadings (scaled for display). The sign of the components was set so that water-stressed genotypes project towards negative PC2 scores in both years. Lower panels, proportion of the variance ($\eta^2$) of PC1 and PC2 explained by genotype, treatment and residual variation. The percentage of total variance explained by each component is given in brackets.

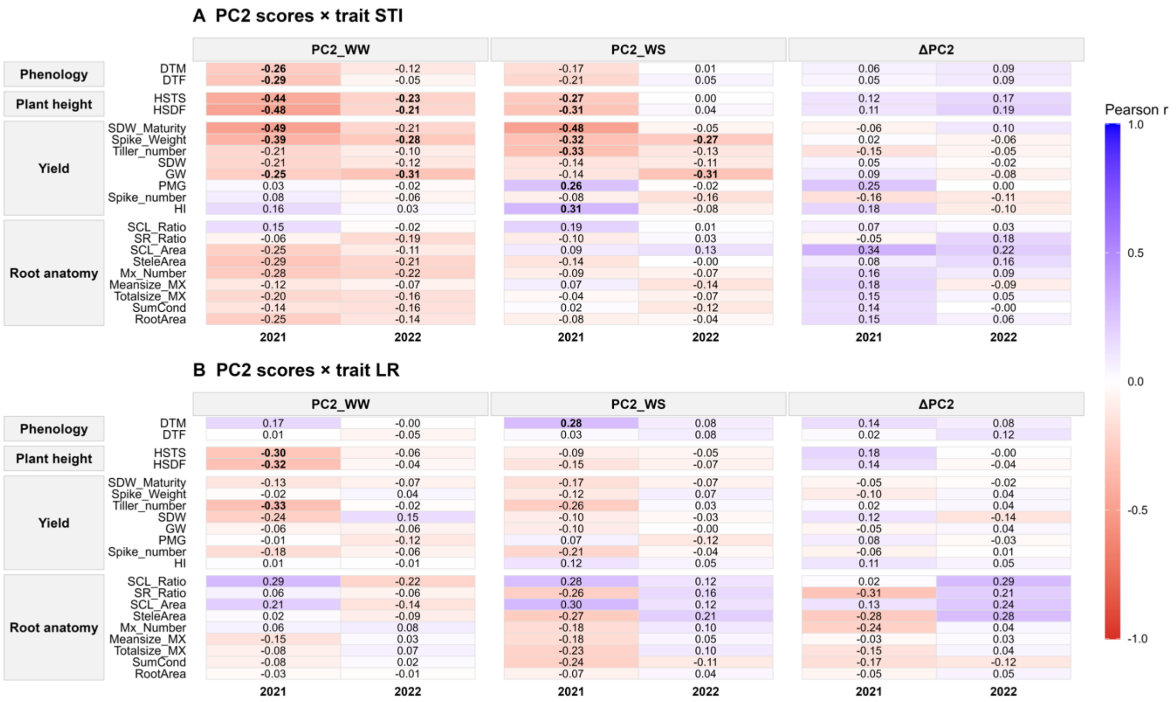


**Fig. S9. Pearson correlations between PC2 scores and agro-morphological traits in each year.** Correlations of PC2 scores under well-watered (PC2_WW) and water-stressed (PC2_WS) conditions, and of their difference (ΔPC2 = PC2_WS − PC2_WW), with (A) stress tolerance indices (STI) and (B) plasticity indices (LR, log2 of the ratio of water-stressed to well-watered values) of phenological, plant height, yield and root anatomical traits. Each cell gives the Pearson r for one year (2021 or 2022). Colour indicates the sign and magnitude of r (blue, positive; red, negative; scale from -1 to 1). Values in bold are significant (FDR-adjusted $p < 0.05$ within trait category). Trait abbreviations are defined in Supplementary Table S1.
